# The Integrator complex terminates promoter-proximal transcription at protein-coding genes

**DOI:** 10.1101/725507

**Authors:** Nathan D. Elrod, Telmo Henriques, Kai-Lieh Huang, Deirdre C. Tatomer, Jeremy E. Wilusz, Eric J. Wagner, Karen Adelman

## Abstract

The transition of RNA polymerase II (Pol II) from initiation to productive elongation is a central, regulated step in metazoan gene expression. At many genes, Pol II pauses stably in early elongation, remaining engaged with the 25-60 nucleotide-long nascent RNA for many minutes while awaiting signals for release into the gene body. However, a number of genes display highly unstable promoter Pol II, suggesting that paused polymerase might dissociate from template DNA at these promoters and release a short, non-productive mRNA. Here, we report that paused Pol II can be actively destabilized by the Integrator complex. Specifically, Integrator utilizes its RNA endonuclease activity to cleave nascent RNA and drive termination of paused Pol II. These findings uncover a previously unappreciated mechanism of metazoan gene repression, akin to bacterial transcription attenuation, wherein promoter-proximal Pol II is prevented from entering productive elongation through factor-regulated termination.

**Highlights:**

- The Integrator complex inhibits transcription elongation at ∼15% of mRNA genes
- Integrator targets promoter-proximally paused Pol II for termination
- The RNA endonuclease of Integrator subunit 11 is critical for gene attenuation
- Integrator-repressed genes are enriched in signaling and growth-responsive pathways

## INTRODUCTION

Dysregulated gene activity underlies a majority of developmental defects and many diseases including cancer, immune and neurological disorders. Accordingly, the transcription of protein-coding messenger RNA (mRNA) is tightly controlled in metazoan cells, and can be regulated at the steps of initiation, elongation or termination. During initiation, transcription factors (TFs) cooperate with coactivators such as Mediator to recruit the general transcription machinery and Pol II to a gene promoter. The polymerase then initiates RNA synthesis and moves downstream from the transcription start site (TSS) into the promoter-proximal region. However, after generating a short, 25-60 nt-long RNA, Pol II pauses in early elongation (Adelman and Lis, 2012). Pausing by Pol II is manifested by the DSIF and NELF complexes, which collaborate to stabilize the paused conformation (Core and Adelman, 2019; Henriques et al., 2013; Vos et al., 2018). Release of paused Pol II into productive elongation requires the kinase P-TEFb, which phosphorylates DSIF, NELF and the Pol II C-terminal domain (CTD), removing NELF from the elongation complex and allowing Pol II to resume transcription into the gene body, with enhanced elongation efficiency (Peterlin and Price, 2006).

Release of paused Pol II into productive RNA synthesis is essential for formation of a mature, functional mRNA. If promoter-paused Pol II becomes permanently arrested or dissociates from the DNA through premature termination, then the process of gene expression is short-circuited, and the gene will not be expressed. Thus, the stability and fate of paused Pol II at a given promoter will have profound effects on gene output. Interestingly, work from a number of laboratories has highlighted that the stability of paused Pol II can differ substantially among genes (Buckley et al., 2014; Chen et al., 2015; Erickson et al., 2018; Henriques et al., 2013; Krebs et al., 2017; Shao and Zeitlinger, 2017). In particular, recent studies of paused Pol II in *Drosophila* revealed a surprising diversity of behaviors following treatment of cells with Triptolide (Trp), an inhibitor of TFIIH that prevents new transcription initiation (Henriques et al., 2018; Krebs et al., 2017; Shao and Zeitlinger, 2017; Vispé et al., 2009). At ∼20% of genes, inhibition of transcription initiation with Trp caused a dramatic reduction of promoter Pol II levels within <2.5 minutes (Henriques et al., 2018). Thus, these genes consistently require new transcription initiation in order to maintain appropriate levels of promoter Pol II. As such, it has been proposed that Pol II undergoes multiple iterative cycles of initiation, early elongation and premature termination at these genes, each time releasing a short, non-functional RNA (Erickson et al., 2018; Kamieniarz-Gdula and Proudfoot, 2019; Krebs et al., 2017; Nilson et al., 2017; Steurer et al., 2018). In contrast, a majority of genes were found to harbor a more stable Pol II, with paused polymerase levels persisting after Trp treatment. In fact, after inhibiting transcription initiation, the median half-life of paused Pol II was ∼10 minutes in both mouse and *Drosophila* systems (Chen et al., 2015; Henriques et al., 2018; Jonkers et al., 2014; Shao and Zeitlinger, 2017). Critically, the distinct stabilities of Pol II observed at different promoters suggests that the lifetime of paused polymerase is modulated to tune gene expression levels. However, the factors that mediate this regulation have yet to be elucidated.

Regulation of promoter-proximal termination is well-described in bacteria, where it is termed *attenuation* (Yanofsky, 1981). Attenuation serves to tightly repress gene activity, even under conditions where the polymerase is recruited to a promoter and initiates RNA synthesis at high levels. Mechanistically, bacterial attenuation often involves destabilization of the RNA-DNA hybrid within the polymerase through RNA structures and/or termination factors with RNA helicase activity (Gollnick and Babitzke, 2002; Henkin and Yanofsky, 2002; Yanofsky, 1981). Similar termination mechanisms are recognized in the yeast *Saccharomyces cerevisiae,* where the Nrd1-Nab3-Sen1 (NNS) complex directs termination using coordinated RNA binding and helicase activities (Bresson and Tollervey, 2018). Intriguingly, the NNS complex, which predominantly drives termination of non-coding RNAs, has also been implicated in premature termination at select mRNA loci (Merran and Corden, 2017; Porrua and Libri, 2015; Sohrabi-Jahromi et al., 2019). However, despite the regulatory potential of promoter-proximal attenuation, a similar phenomenon has not yet been described in metazoan cells. In particular, it remains unclear whether higher eukaryotes possess a termination machinery that promotes dissociation of paused early elongation complexes.

Elongating Pol II is typically extremely stable, with formation of a mature mRNA often involving transcription of many kilobases without Pol II dissociation from DNA. Termination at mRNA 3’-ends involves recognition of specific sequences by cleavage and polyadenylation (CPA) factors, and slowing of Pol II elongation. CPSF73, a component of the CPA complex, utilizes a β-lactamase/β-CASP domain (Mandel et al., 2006) to cleave pre-mRNA, producing both a substrate for polyadenylation and a free 5’ end on the nascent RNA still engaged with Pol II. This 5’ end lacks the protective 7-methy-G cap, allowing it to be targeted by the Xrn2 exonuclease, which ultimately leads to termination (Eaton et al., 2018). Hence, cleavage of the nascent RNA is coupled to the termination of elongation and dissociation of Pol II from template DNA, as well as degradation of the associated short RNA. Although the CPA machinery typically functions at gene 3’ ends, there are examples of premature cleavage and polyadenylation (PCPA) occurring within gene bodies, especially within intronic regions (Kamieniarz-Gdula and Proudfoot, 2019; Venters et al., 2019). However, whether this machinery is involved in RNA cleavage and termination of promoter proximal Pol II remains unknown.

We set out to determine the causes of differential stability of paused Pol II across mRNA genes. In particular, we were interested in defining factors that might render promoter Pol II susceptible to premature termination and the release of short, immature RNAs (Erickson et al., 2018; Henriques et al., 2013; Krebs et al., 2017; Nilson et al., 2017; Shao and Zeitlinger, 2017; Steurer et al., 2018). Strikingly, we discovered that the Integrator complex is enriched at mRNA promoters with unstable Pol II pausing. The 14-subunit, metazoan-specific, Integrator complex was initially reported to be exclusively required for cleavage and 3’-end formation of small nuclear RNAs (snRNAs) involved in splicing (Baillat et al., 2005). However, subsequent work has suggested a broader role, including at signal-responsive mammalian genes (Gardini et al., 2014; Lai et al., 2015; Skaar et al., 2015; Stadelmayer et al., 2014). Our work elucidates this role and reveals that Integrator targets paused Pol II at selected protein-coding genes and enhancers, to mediate premature termination. Notably, the Integrator complex, like the CPA machinery, possesses an RNA endonuclease, and we find that this activity is critical for gene repression. Thus, our findings unearth transcription attenuation as a conserved, broad mode of gene control in metazoan cells.

## RESULTS

The underlying cause for the short lifetime of paused Pol II at a subset (∼20%) of *Drosophila* protein coding genes is not understood (Buckley et al., 2014; Henriques et al., 2018; Krebs et al., 2017; Shao and Zeitlinger, 2017). One potential explanation for the brief lifetime of Pol II near these promoters is that paused polymerase is quickly released into productive elongation. This model would predict that such genes would generally have lower levels of Pol II near their promoters, and more Pol II elongating within gene bodies. An alternative possibility is that fast Pol II turnover at these genes results from rapid transcription termination of promoter-paused Pol II. The key prediction of this latter model is that these genes would display lower levels of productively elongating Pol II within gene bodies.

To evaluate these possibilities, we compared nascent RNA profiles determined by PRO-seq, a single-nucleotide resolution method for mapping active and transcriptionally engaged Pol II (Kwak et al., 2013). Genes were stratified into four clusters based on their Pol II decay rate following Trp treatment (Henriques et al., 2018; Krebs et al., 2017) and were analyzed for PRO-seq signals near the promoter or within the gene body. We found that genes with short-lived promoter Pol II occupancy (defined as half-life upon Trp-treatment <2.5 min) have significantly lower elongating Pol II levels than other gene classes (Figures 1A and S1A), despite modestly higher promoter Pol II signals. These data are thus consistent with a model wherein Pol II is efficiently recruited to these promoters, but fails to enter productive elongation, possibly due to premature termination (Krebs et al., 2017).

**Figure 1.**
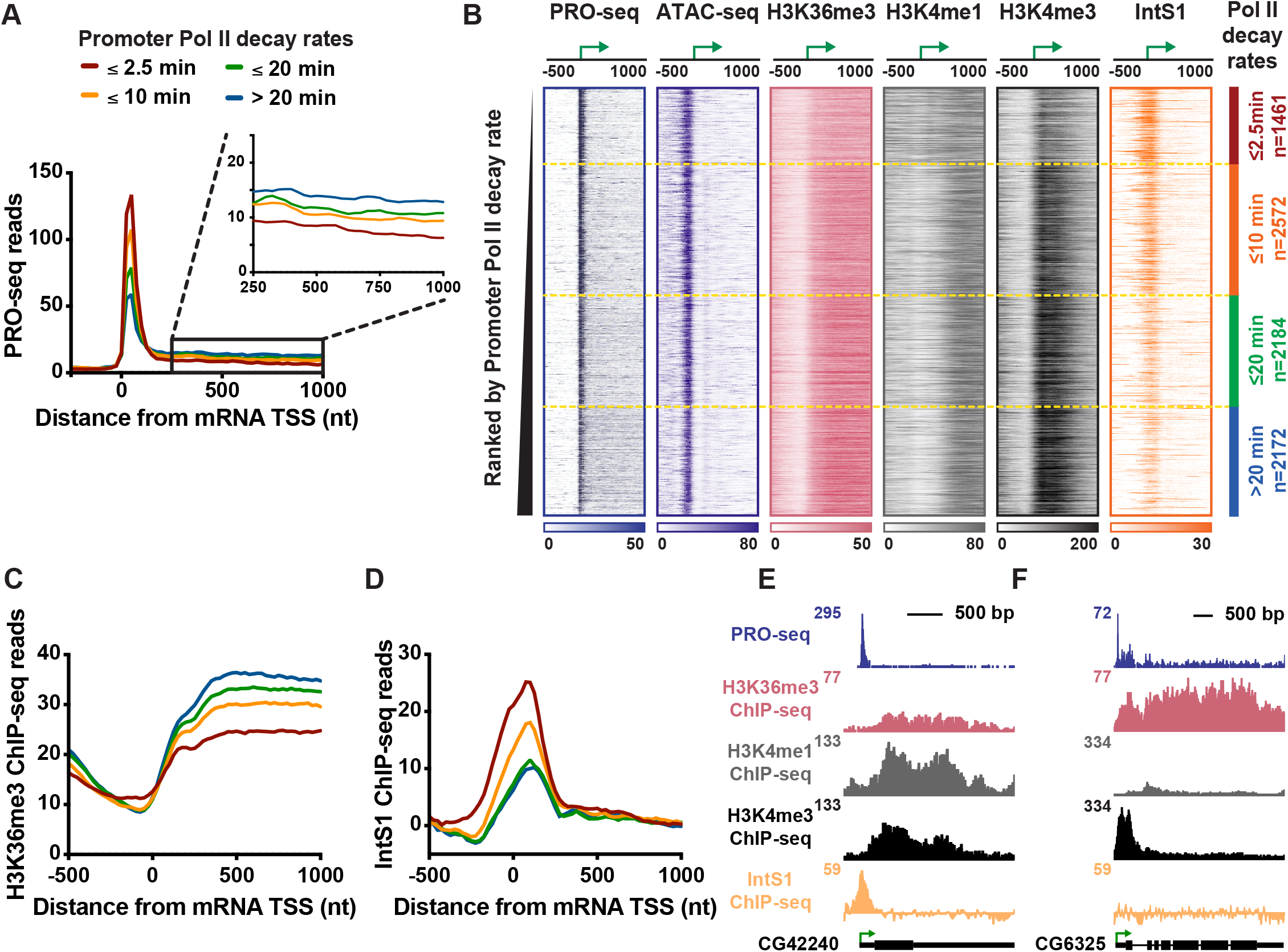
Genes with highly unstable promoter Pol II are characterized by poor transcription elongation and enriched binding of Integrator. (A) The average distribution of PRO-seq signal is shown at mRNA transcription start sites (TSSs), with genes divided into four groups based on Pol II promoter decay rates following Triptolide treatment (groups defined in Henriques *et al*., 2018). Inset shows the gene body region. Read counts are summed in 25-nt bins. (B) Heatmap representations of PRO-seq and ATAC-seq signal, along with ChIP-seq reads for H3K36me3, H3K4me1 and H3K4me3 histone modifications and the Integrator subunit 1 (IntS1). Data are aligned around mRNA TSSs, shown as a green arrow (n=8389). Data are ranked by Promoter Pol II decay rate, where promoters with fastest decay rates (≤2.5 min) are on top. Dotted line separates each group of genes. (C and D) Average distribution of (C) H3K36me3, and (D) IntS1, ChIP-seq signal is shown, aligned around TSSs and divided into groups based on Pol II decay rate, as in A. (E and F) Example gene loci, representative of genes in the (E) fast, or (F) slow, Pol II promoter decay groups, displaying profiles of PRO-seq and ChIP-seq signals, as indicated. See also Figure S1.

To evaluate this prediction and define factors that might contribute to this behavior, we computationally assessed a comprehensive repertoire of ChIP-seq data (Baumann and Gilmour, 2017; Henriques et al., 2018; Ho et al., 2014; Kaye et al., 2018; modENCODE Consortium et al., 2010; Weber et al., 2014). Specifically, we sought to identify factors enriched (or de-enriched) at gene promoters where pausing is unstable as compared to other promoters (see Methods). Chromatin accessibility was observed to be consistent across Pol II decay classes (as assessed by ATAC-seq, Figures 1B and S1B), consistent with the similar promoter Pol II levels observed. However, reduced levels of tri-methylated H3 Lysine 36 (H3K36me3) were noted within genes harboring unstable promoter Pol II (Figures 1C and S1C). The H3K36me3 mark is deposited during productive elongation, and H3K36me3 levels typically correlate with transcription activity (Venkatesh and Workman, 2015; Wagner and Carpenter, 2012). Thus, the observed, low H3K36me3 signal indicates weak transcription elongation at genes with unstable Pol II, consistent with PRO-seq data. Conversely, genes with stable pausing exhibited stronger transcription activity and higher levels of H3K36me3 (Figures 1A and 1B), in agreement with recent work (Tettey et al., 2019).

Genes with unstable Pol II also displayed a significant enrichment in H3K4 mono-methylation (H3K4me1) and lower tri-methylation of H3K4 (H3K4me3) and as compared to genes with more stable pausing (Figures 1B, S1D and S1E). This finding suggests that H3K4 methylation levels increase near promoters as Pol II stability and residence time increases, in agreement with a recent study in yeast (Soares et al., 2017). Intriguingly, elevated H3K4me1 levels, with deficiencies in H3K36me3, H3K4me3 and productive RNA elongation are considered to be characteristics of enhancers (ENCODE Project Consortium, 2012; Kim and Shiekhattar, 2015; Perissi et al., 2010). Enhancers are also characterized by unstable Pol II and the production of short RNAs (Henriques et al., 2018), suggesting a connection between the chromatin signatures typical of enhancers and defective or inefficient transcription elongation.

To define additional factors that could contribute to the transcriptional properties of these genes, we analyzed ChIP-seq profiles of non-chromatin proteins. We found the Integrator subunit 1 (IntS1) among the most significantly enriched factors at genes with unstable Pol II (Figures 1D, 1E and S1F). This is an interesting finding, given that Integrator is implicated in the biogenesis of enhancer-derived RNAs (eRNAs) in human cells (Lai et al., 2015), and further underscores the similarity between this class of genes and enhancers. To confirm these results, we conducted ChIP-seq using an antibody raised against another *Drosophila* Integrator subunit, IntS12, and found a highly similar enrichment at genes with unstable Pol II (Figure S1G).

In summary, genes with unstable promoter Pol II display typical levels of Pol II recruitment and promoter DNA accessibility, but significantly diminished Pol II elongation. These genes display chromatin features reminiscent of enhancers, suggestive that a lack of stable pausing and transcription elongation has considerable consequences on local chromatin modifications (Figures 1E and 1F). Interestingly, these genes also show elevated occupancy by Integrator, a factor known to mediate RNA cleavage and Pol II termination at non-coding RNA loci.

### Loss of Integrator leads to reduced promoter-proximal termination and upregulation of gene expression

Two Integrator subunits, IntS11 and IntS9, are paralogs of the CPA proteins CPSF73 and CPSF100, respectively. IntS11, like CPSF73, has a β-lactamase/β-CASP domain and harbors endonuclease activity. Moreover, similar to CPSF73/100, IntS11 forms a heterodimer with IntS9 and this association is essential for function (Wu et al., 2017). This similarity suggests that Integrator might be capable of mediating transcription termination at protein-coding genes using a mechanism related to that of the CPA machinery. To evaluate this possibility, IntS9 was depleted using RNA interference (RNAi) for 60 hours (Figure S2A), followed by polyA-selected RNA-seq to identify mRNA expression changes. Consistent with the reported stability of snRNAs, their steady-state levels were not perturbed during the relatively short time course of RNAi (Figure S2B), and very few differences in splicing events were observed in IntS9-depleted cells (see Methods). Thus, short-term loss of Integrator has minimal effects on snRNA functionality or splicing patterns. Nonetheless, genes with any evidence of altered splicing in IntS9-depleted cells were removed from all further analyses, enabling us to solely focus on transcriptional targets of Integrator.

Our analysis revealed 723 upregulated and 163 downregulated mRNAs upon IntS9 depletion (Figure 2A), suggesting that *Drosophila* Integrator is predominantly a transcriptional repressor. The expression changes observed upon IntS9 RNAi were validated using RT-qPCR at selected genes (Figure S2C). Gene Ontology analysis of upregulated transcripts shows significant enrichment in signal-responsive pathways, including metabolic, receptor and oxidoreductase activities, as well as Epidermal Growth Factor (EGF)-like protein domains (Figure S2D). Consistently, work on mammalian Integrator has implicated this complex in EGF-responsive gene activity (Gardini et al., 2014).

**Figure 2.**
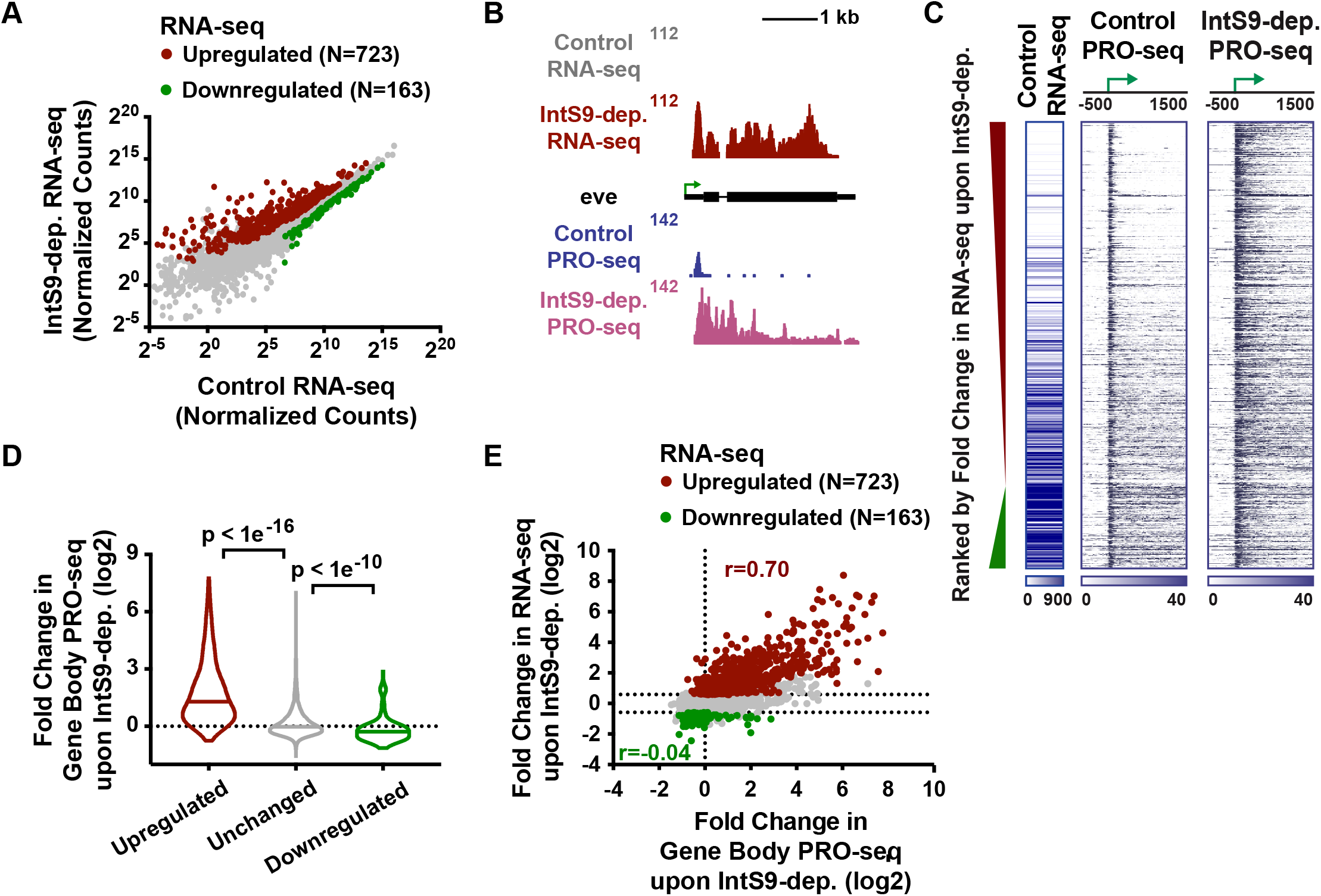
The Integrator complex attenuates expression of protein-coding genes. (A) Drosophila cells were treated for 60 h with control dsRNA, or dsRNA targeting IntS9 (N=3). Normalized RNA-seq signal is shown, with significantly affected genes defined as P<0.0001 and fold change >1.5. (B) even skipped (eve, CG2328) locus displaying profiles of RNA-seq and PRO-seq in control and IntS9-depleted cells. (C) Heatmap representations of RNA-seq levels are shown, along with PRO-seq reads from control and IntS9-depleted cells (treated as in A). The location of mRNA TSSs is indicated by an arrow. Genes that are upregulated or downregulated upon IntS9-depletion in RNA-seq are shown, ranked from most upregulated to most downregulated. (D) Violin plots depict the change in gene body PRO-seq signal upon IntS9-depletion for each group of genes. IntS9-affected genes are defined as in A, as compared to 8613 unchanged genes. Plots show the range of values, with a line indicating median. P-values are calculated using a Mann-Whitney test. (E) Comparison of fold changes in RNA-seq and PRO-seq signals upon IntS9-depletion is shown. Pearson correlations are shown separately for upregulated and downregulated genes, indicating good agreement between steady-state RNA-seq and nascent PRO-seq signals for upregulated genes, but little correspondence for downregulated genes. See also Figure S2.

To probe the mechanisms by which Integrator regulates gene expression, we directly monitored nascent RNA synthesis using PRO-seq in control or IntS9-depleted cells. Critically, PRO-seq is amenable to spike-in normalization, allowing us to ensure that quantitative differences between samples can be accurately measured (Methods). PRO-seq in control cells revealed that Pol II is effectively recruited to IntS9-repressed promoters, but the polymerase often fails to transition into productive elongation (Figures 2B and 2C). In fact, genes upregulated upon IntS9 depletion exhibited significantly higher PRO-seq signal at promoters, yet lower PRO-seq signal within gene bodies and lower mRNA expression than unaffected genes (Figure S2F). These data demonstrate that Integrator does not repress transcription initiation but rather prevents the transition of promoter-proximal Pol II into productive RNA synthesis, perhaps by mediating transcription termination. Consistent with this possibility, depletion of IntS9 relieved the strong block to productive elongation at upregulated genes, allowing a significant, median 3-fold, increase of PRO-seq signal within gene bodies (Figures 2C and 2D).

There was highly significant overlap between transcripts deemed significantly upregulated in PRO-seq and RNA-seq experiments, confirming that the upregulated mRNA production observed upon IntS9 depletion generally results from increased transcription elongation at these genes (Figures 2E and S2G). In contrast, decreases in RNA-seq signal were not well-reflected in PRO-seq levels, with fold-changes between the assays correlating poorly (Figure 2E). Indeed, only 29 transcripts were defined as downregulated by IntS9-depletion in both the RNA-seq and PRO-seq assays (Figure S2G). We thus conclude that the dominant transcriptional effect of *Drosophila* Integrator at protein-coding genes is in transcription repression.

### The Integrator RNA endonuclease is required for transcriptional repression

The above data suggest that Integrator might use its endonuclease activity to catalyze transcription termination of paused Pol II. To test this model, and determine whether IntS11 catalytic function is required for gene repression, we took advantage of a previously described mutant (IntS11 E203Q; Figure 3A) that abrogates endonuclease function yet retains the integrity of the Integrator complex (Baillat et al., 2005). We treated *Drosophila* cells for 60 hours with either control RNAi or with RNAi targeting the IntS11 UTRs and also re-expressed either wild-type IntS11 or the E203Q mutant in cells depleted of endogenous IntS11 (Figure S3A). RNA from these cells was isolated and subjected to poly(A)-enriched RNA-seq. As with IntS9 depletion, mature snRNA levels are not perturbed by IntS11 knockdown, and the major effect was upregulation of transcription (Figures S3B and S3C). Further, the levels of gene upregulation observed upon depletion of IntS9 or IntS11 were highly concordant (Figures 3B and S3C). In contrast, there was less agreement and smaller effect sizes observed at downregulated genes (Figures 3B and S3C).

**Figure 3.**
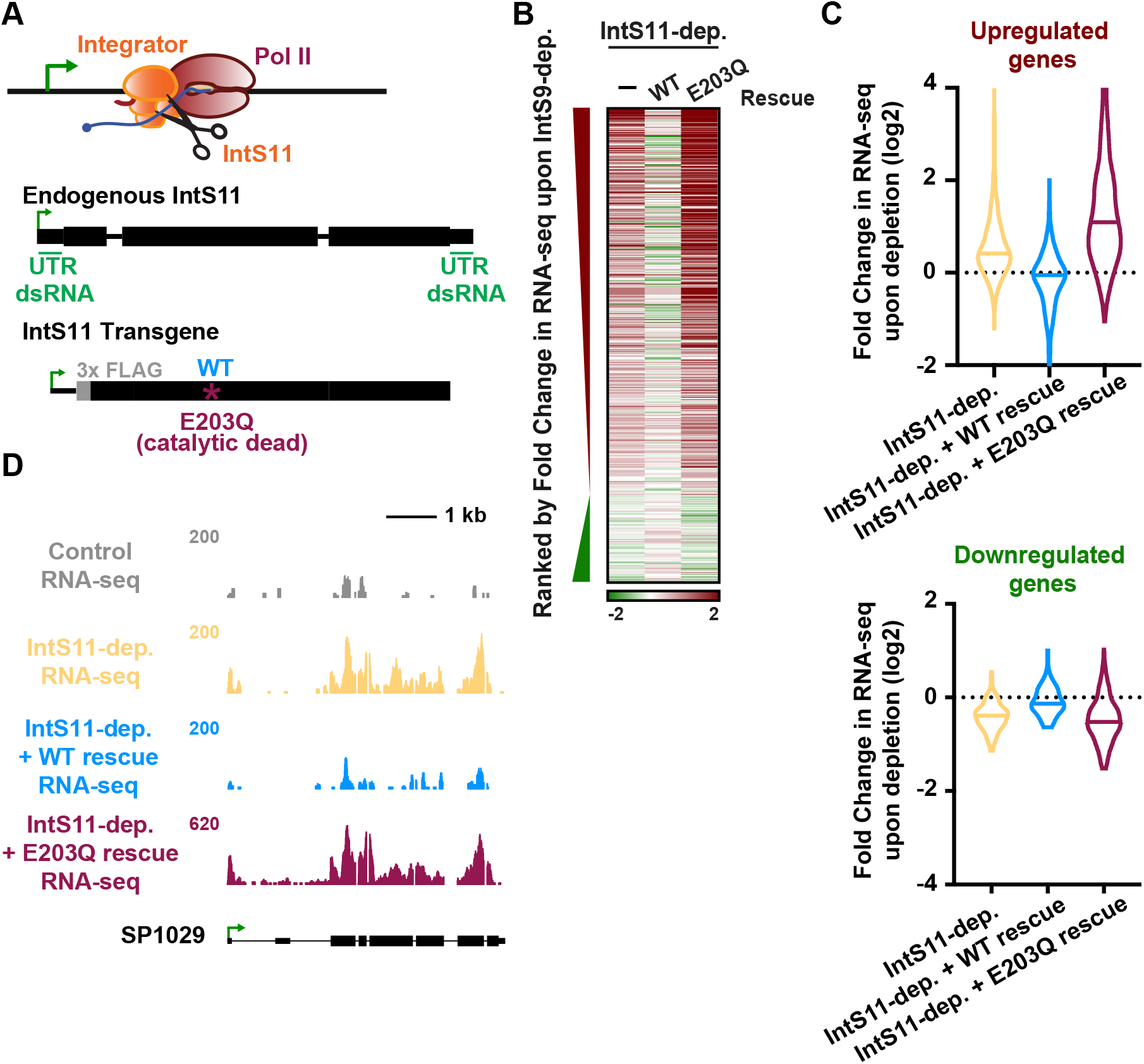
Integrator subunit 11 (IntS11) endonuclease activity is essential for altered protein-coding gene expression. (A) The IntS11 subunit of Integrator harbors RNA endonuclease activity (depicted as scissors). To test the importance of this activity, cells were depleted of IntS11 and rescued using a stably integrated transgene expressing WT IntS11, or IntS11 with a mutation that disrupts endonuclease activity (E203Q). To specifically deplete endogenous IntS11 from the rescue cell lines, a dsRNA targeting the untranslated (UTR) regions of endogenous IntS11 (green) was used. Cells were treated for 60 h with control or IntS11 UTR RNAi (N=3, see Methods), and RNA harvested for RNA-seq. (B) Heatmap representations of RNA-seq fold changes in IntS11-depleted cells, as compared to cells rescued with WT or E203Q mutant. Genes shown are those affected upon IntS9-depletion, ranked by fold-change as in Figure 2C. (C) Fold Change in RNA-seq signal upon IntS11-depletion at genes (top) upregulated (N=723) or (bottom) downregulated by IntS9-depletion (N=163). Changes in RNA-seq levels as compared to the parental cell line are shown in IntS11-depleted cells, and those rescued by WT or E203Q mutant IntS11. Violin plots show range of values, with a line indicating median. (D) SP1029 (CG11956) locus showing an upregulated gene whose expression is rescued by WT IntS11, but not by the catalytic dead mutant (E203Q mutation). RNA-seq tracks are shown in control cells and each of the treatments. See also Figure S3.

The vast majority of gene expression changes observed in IntS11-depleted cells were restored to normal, control levels upon expression of the wild-type IntS11 (Figures 3B, 3C and S3D). In contrast, expression of the E203Q mutant not only failed to rescue the IntS11 depletion but exacerbated the knockdown phenotype, supportive of a dominant negative effect of the catalytically inactive IntS11 protein (Figures 3B, 3C and S3D). The results observed by RNA-seq (e.g. Figure 3D) were confirmed by RT-qPCR (Figure S3E). Together, these data indicate that depletion of either IntS9 or IntS11 lead to alteration of a similar set of protein-coding genes and that the IntS11 endonuclease activity is essential for the function of Integrator at these loci.

### Integrator attenuates mRNA transcription

The critical involvement of the IntS11 endonuclease in gene repression by Integrator supports a model wherein RNA cleavage triggers premature termination. To further evaluate this model, we defined the full repertoire of transcriptional targets of Integrator, by comparing spike normalized PRO-seq signals in gene bodies between control and IntS9-depleted samples (see Methods). We found 1204 transcripts with significantly more elongating Pol II upon depletion of Integrator (Figure 4A), and 210 with reduced gene-body Pol II signal. This reveals that transcription of ∼15% of active *Drosophila* genes is upregulated upon loss of Integrator activity.

**Figure 4.**
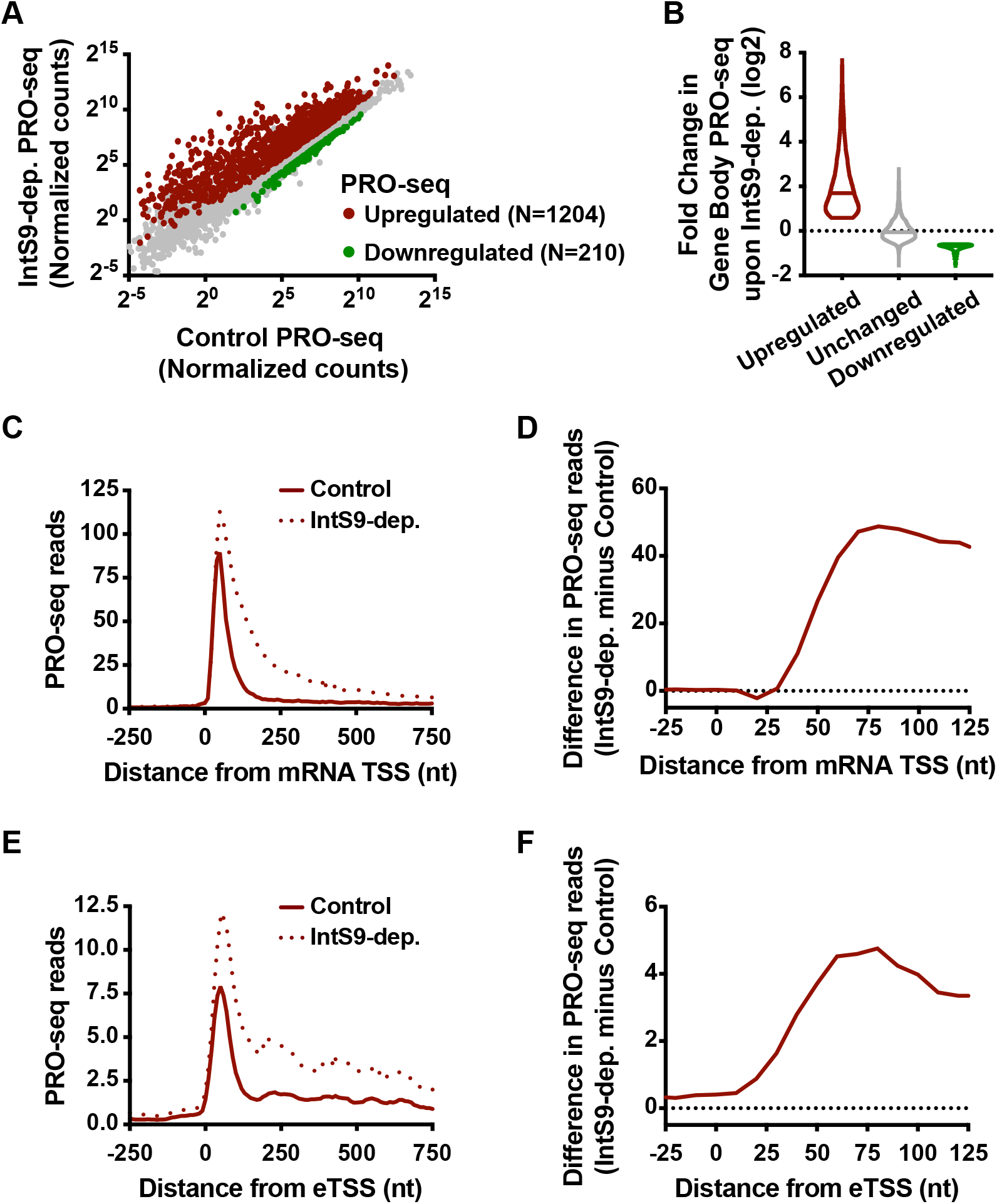
Integrator represses productive elongation by Pol II at genes and enhancers. (A) Drosophila cells were treated for 60 h with control or IntS9 RNAi (N=3). Normalized PRO-seq signal across gene bodies is shown, with IntS9-affected genes defined as P<0.0001 and fold change >1.5. (B) Violin plots depict the change in gene body PRO-seq signal upon IntS9-depletion for each group of genes. IntS9-affected genes are defined as in A, as compared to unchanged genes (N=8085). Violin plots show range of values, with a line indicating median. (C) Average distribution of PRO-seq signal in control and IntS9-depleted cells is shown at upregulated genes. (D) The difference in PRO-seq signal between IntS9-depleted and control cells for upregulated genes is shown. Increased signal in IntS9-depleted cells is consistent with the position of Pol II pausing, from +25 to +60 nt downstream of the TSS. (E) Average distribution of PRO-seq reads from control and IntS9-depleted cells are displayed, centered on enhancer transcription start sites (eTSS) that are upregulated upon IntS9 RNAi (N=228). (F) Difference in PRO-seq signal between IntS9-depleted and control cells for IntS9-upregulated enhancer RNAs. Note that signal increases at enhancers in the same interval (+25-60 nt from TSS) as at coding loci. See also Figure S4.

Gene ontology analyses of the genes upregulated in PRO-seq agreed well with those from RNA-seq, highlighting metabolic, oxidoreductase and EGF pathways (Figure S4A and S2D). In contrast, enriched pathways for the downregulated genes in PRO-seq overlapped little with those enriched among RNA-seq downregulated genes (Figures S4B and S2E), in agreement with the lack of concordance between nascent transcription and steady-state RNA levels within the downregulated gene sets (Figures 2E and S2G; only 29 genes downregulated in both PRO-seq and RNA-seq). Thus, we focused our attention on the much larger set of upregulated loci. The increase in gene body PRO-seq signal upon IntS9-depletion was substantial at upregulated genes, with a median increase of over 3.3-fold (Figure 4B). As anticipated, the majority of this increase in actively engaged Pol II is evident in PRO-seq signal near TSSs (Figure S4C). Thus, we conclude that Integrator typically acts on promoter-proximal Pol II, and that loss of Integrator results in increased levels of engaged polymerase that successfully transition from promoter regions into productive elongation.

We then wished to distinguish between models wherein Integrator catalyzes promoter-proximal termination vs. those wherein Integrator prevents escape of promoter-associated Pol II into productive elongation. We evaluated the PRO-seq signal at genes upregulated upon depletion of IntS9. If Integrator holds Pol II near promoters, then IntS9 depletion should release this paused Pol II into gene bodies, resulting in less promoter-proximal PRO-seq signal and an increase in signal downstream. In contrast, if Integrator stimulates termination and dissociation of paused Pol II, then IntS9 depletion should increase PRO-seq signals both promoter-proximally and within genes. In support of a termination model, we observed that IntS9 depletion resulted in increased PRO-seq signal near promoters, as well as in gene bodies (Figure 4C). Strikingly, the increase in PRO-seq signal from IntS9-depleted cells localized precisely at the position of Pol II pausing, in the window from 25-60 nt into the gene (Figure 4D). This finding supports that Integrator targets promoter-paused Pol II and prevents its transition into productive RNA synthesis, likely through premature termination.

To determine whether Integrator similarly targets paused Pol II at enhancers, we made use of a comprehensive set of *Drosophila* enhancer transcription start sites (eTSSs) we recently defined (Henriques et al., 2018). We note that these sites were rigorously defined both functionally, in plasmid-based enhancer reporter assays (Arnold et al., 2013; Zabidi et al., 2015) and spatially, with the TSSs of enhancer RNAs (eRNAs) mapped at single-nucleotide resolution (Henriques et al., 2018). This dataset thus allows for a high-resolution analysis of Integrator activity at functionally-confirmed, transcriptionally active enhancer loci at the genome-level. We focused on 1498 intergenic eTSSs, to avoid confounding signals from enhancers within annotated genes, and defined differentially transcribed loci using PRO-seq data as we had for mRNA genes (see Methods). We observed increased transcription at ∼15% of eTSSs in IntS9-depleted cells (N=228), a similar fraction to mRNAs (Figure S4D) and find only 38 eTSSs with downregulated transcription. Thus, at enhancers, like at protein-coding genes, Integrator plays a generally repressive role in transcription elongation, and targets only selected loci. Importantly, many eRNA loci are not affected by loss of Integrator (Figure S4E), consistent with work implicating CPA and other machineries in eRNA 3’ end formation (Austenaa et al., 2015; Ogami et al., 2017).

The parallel in the behavior of Integrator at protein-coding and non-coding loci is further emphasized by the profile of PRO-seq at upregulated eTSSs (compare Figures 4E and 4C), where loss of Integrator causes an increase of PRO-seq signal precisely in the region of Pol II pausing (compare Figures 4F and 4D). We conclude that the function of Integrator is highly similar at coding and non-coding RNA loci: a comparable subset of TSSs are affected by Integrator, and Integrator depletion causes increased Pol II near TSSs and higher levels of release downstream into productive elongation.

### Integrator is widely associated with mRNA promoter regions

The mechanism for Integrator-mediated 3’ end formation at snRNA loci involves both selective recruitment of Integrator to snRNA promoters and recognition of a degenerate motif near snRNA 3’ ends that promotes IntS11 cleavage activity (Baillat and Wagner, 2015; Hernandez, 1985; Hernandez and Weiner, 1986). Interestingly, several factors implicated in recruiting Integrator to snRNA genes are also found at protein coding loci, such as the pause-inducing factors DSIF and NELF (Stadelmayer et al., 2014; Yamamoto et al., 2014), and phosphorylation on the Pol II C-terminal domain (CTD) repeats at Serine 7 residues (Egloff et al., 2007; Kim et al., 2010). Consistent with this, Integrator has been observed to associate with some mRNA promoters in human systems (Gardini et al., 2014; Skaar et al., 2015; Stadelmayer et al., 2014). However, it has not been fully explored how well the localization of Integrator at promoters corresponds to its gene regulatory activities at a genome-wide level.

To address this question, we investigated the global localization of Integrator using our ChIP-seq datasets. We find that IntS1 and IntS12 subunits showed highly correlated localization across snRNA (r=0.99) and mRNA promoters (r=0.89) (Figures S5A and S5B), with a strong enrichment near mRNA transcription start sites (Figures 5A and S5C). However, Integrator signal at promoters correlated only weakly with levels of paused Pol II as determined by promoter PRO-seq signal (Figure S5B, r=0.39). Whereas these findings are consistent with Pol II, DSIF and NELF representing interaction surfaces for Integrator, they also indicate that association of Integrator with mRNA promoters is not strictly tied to paused Pol II levels. We thus asked whether there was enrichment of IntS1 or IntS12 occupancy at genes that are upregulated upon depletion of IntS9. Indeed, genes repressed by Integrator were significantly enriched in both IntS1 and IntS12 ChIP-seq signal as compared to genes unaffected by Integrator depletion (Figures 5B, 5C, 5D and S5D). In fact, levels of Integrator observed at IntS9-repressed promoters were even higher than levels at snRNAs (Figure 5D and S5G). We noted, however, that Integrator ChIP-seq signals at genes with unchanged expression upon IntS9 RNAi were well above background levels, suggesting that Integrator is also recruited to promoters where it remains inactive.

**Figure 5.**
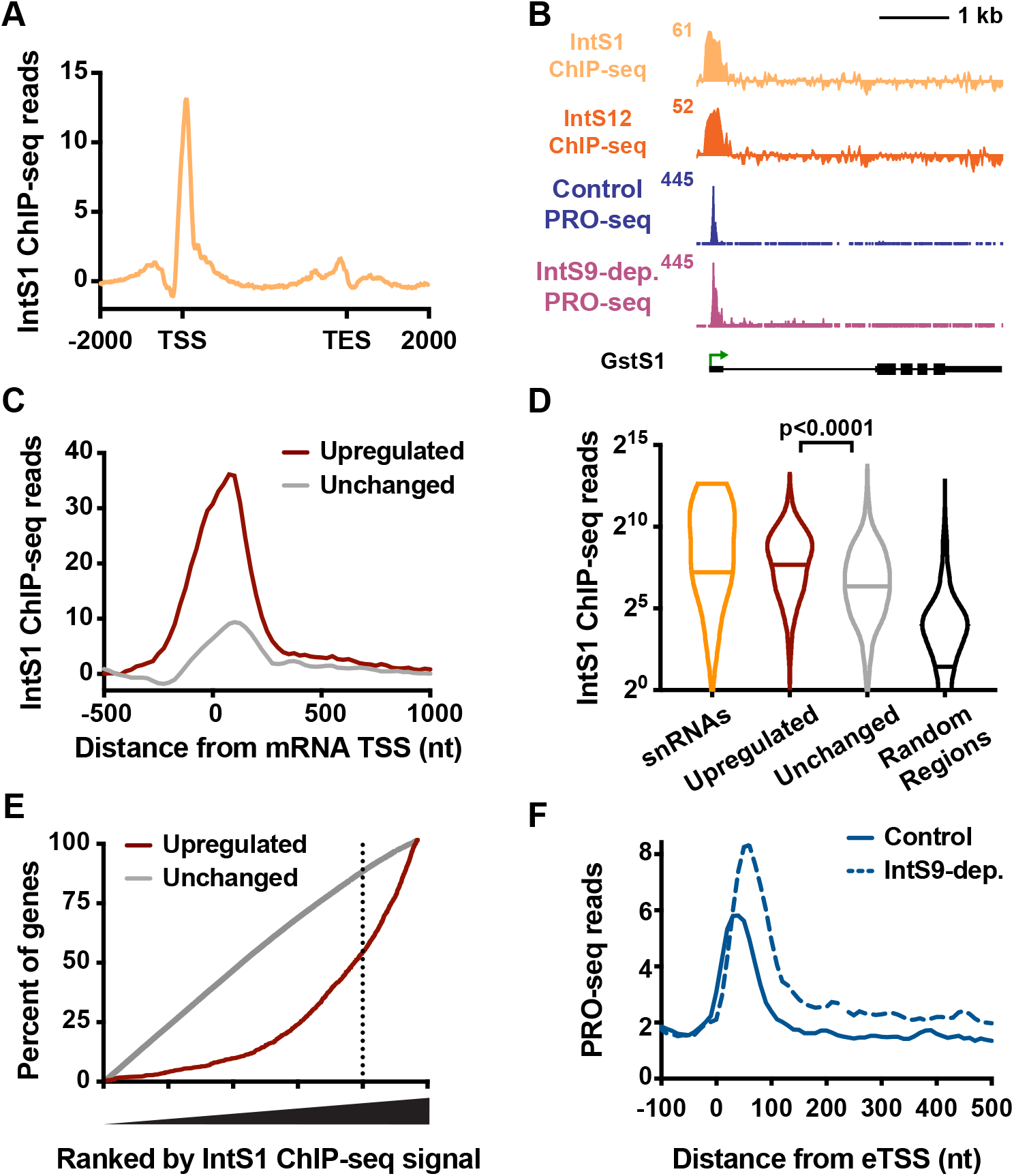
Integrator binding is enriched at promoters of target genes. (A) Distribution of IntS1 ChIP-seq signal along the transcription units of all active mRNA genes (N=9499). Windows are from 2 kb upstream of the TSS to 2 kb downstream of the transcription end site (TES). Bin size within genes is scaled according to gene length. (B) Example locus (GstS1) of an upregulated gene upon IntS9-dep. showing PRO-seq and Integrator ChIP-seq. (C) Metagene analysis of average IntS1 ChIP-seq signal around promoters of upregulated (N=1204) and unchanged (N=8085) mRNA genes in IntS9-depleted cells. Data are shown in 25-nt bins. (D) Promoter-proximal IntS1 ChIP-seq reads for each group of sites: snRNAs (N=31), upregulated or unchanged genes, and randomly-selected intergenic regions (N=5000). Violin plots show range of values, with a line indicating median. P-values are calculated using a Mann-Whitney test. (E) All active genes (N=9499) were rank ordered by increasing IntS1 ChIP-seq signal around promoters (± 250bp), and the cumulative distribution of upregulated or unchanged genes across the range of IntS1 signal is shown. IntS1 levels at unchanged genes show no deviation from the null model, but upregulated genes display a significant bias towards elevated IntS1 ChIP-seq signal. (F) Average distribution of PRO-seq signal at eTSSs bound by the Integrator complex (N=691) in control and IntS9-depleted cells is shown. See also Figure S5.

To further investigate the relationship between Integrator binding and activity, we rank ordered all active mRNA promoters by their IntS1 ChIP-seq signal, and calculated cumulative distributions of Integrator-repressed and unchanged genes across this ranking (Figure 5E). This analysis demonstrated that Integrator exhibits the full spectrum of binding levels at unchanged genes. However, IntS9-repressed genes were clearly and significantly biased towards higher IntS1 occupancy (Figure 5E, >50% of IntS9-repressed genes fall within the top 20% of IntS1 levels, whereas only 15% of unchanged genes fall in this group). Thus, like at the snRNAs, Integrator recruitment to an mRNA promoter is not sufficient to dictate function, but high-level Integrator occupancy is typically associated with activity.

To determine whether increased recruitment of Integrator was also related to functional outcomes at enhancers, we identified eTSSs that exhibited significant peaks of IntS1/IntS12 signal (Figure S5E). Comparing PRO-seq at these loci in control vs. IntS9-depleted conditions demonstrated that Integrator-bound eTSSs showed increased transcription elongation upon IntS9 RNAi (Figure 5F). In contrast, no significant change in PRO-seq signal was observed at Integrator-unbound eTSSs upon depletion of IntS9 (Figure S5F). We conclude that functional mRNA and eRNA targets of Integrator display greater recruitment of this complex. Although the factors governing this elevated recruitment of Integrator at snRNA or other loci remain to be elucidated, our results underscore a common behavior for Integrator at coding and non-coding loci.

### Integrator mediates cleavage of nascent RNA and promoter-proximal termination

Taken together, our results are most consistent with Integrator serving as a promoter-proximal cleavage and termination factor for a set of protein-coding genes. To definitively test this possibility, we investigated the short, TSS-associated RNAs that would accompany Pol II termination. In particular, we used Start-seq (Henriques et al., 2018; Nechaev et al., 2010; Williams et al., 2015) to identify RNAs under 100 nt in length that were 3′ oligoadenylated, a modification that can be detected on a minor fraction of RNAs released by Pol II during termination (Figure 6A). Such oligoadenylated termination products are subject to degradation and normally very short-lived, but are stabilized in cells depleted of the RNA Exosome. Accordingly, following depletion of the Exosome subunit Rrp40, we observed significantly more oligoadenylated short RNAs from IntS9-repressed genes than unchanged genes (Figure 6B). Strikingly, the 3′ ends of these oligoadenylated RNAs are highly and specifically enriched within the region of Pol II pausing (Figure 6C).

**Figure 6.**
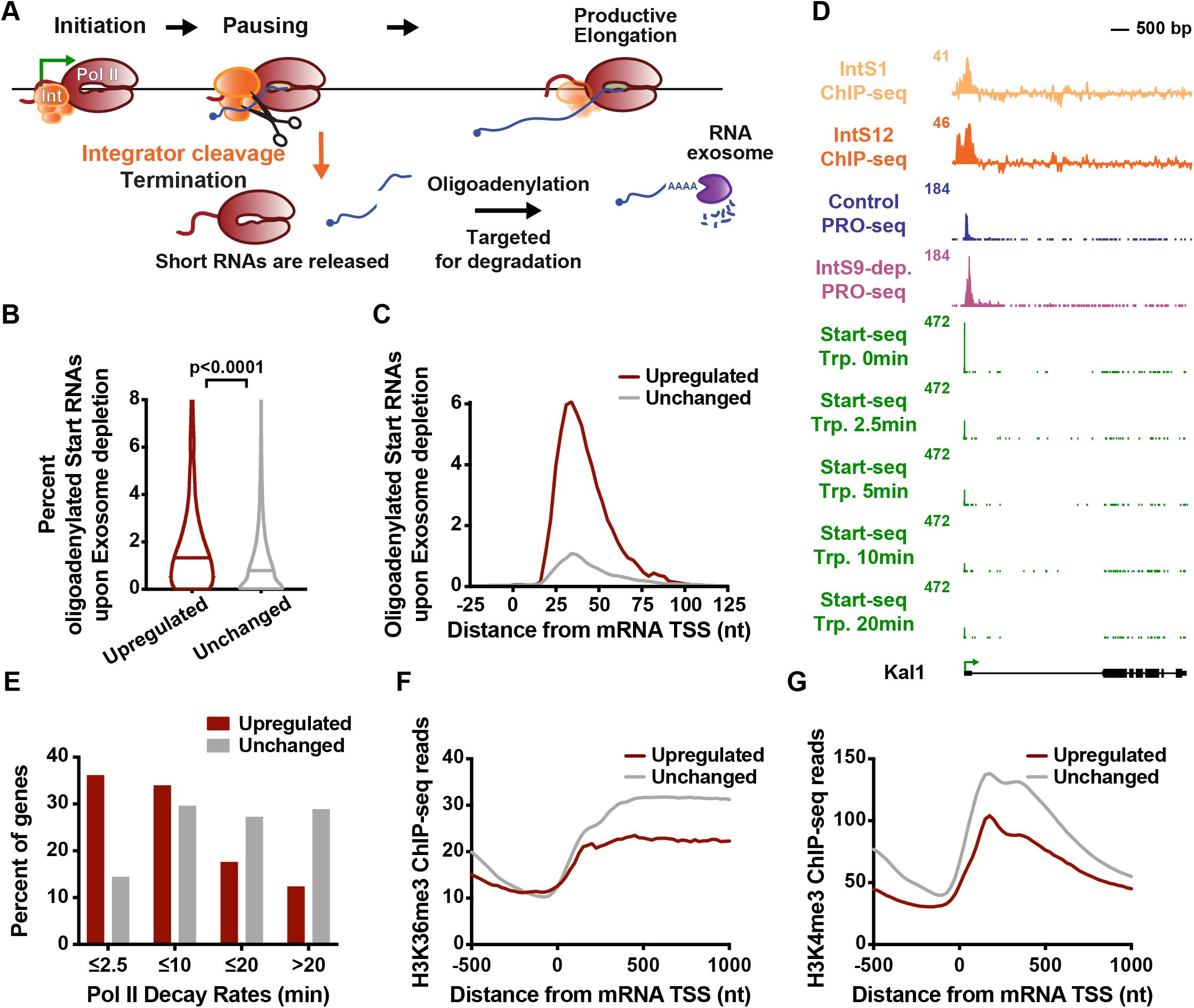
Integrator attenuates mRNA expression through promoter-proximal termination. (A) Schematic of transcription cycle with possible fates of Pol II. Paused Pol II can enter into productive elongation or terminate and release a short RNA. A small fraction of released RNA is oligoadenylated to facilitate degradation by the RNA exosome. (B) The percent of Start RNA reads bearing oligoadenylated 3’ ends in exosome-depleted (Rrp40 subunit) cells is shown for each gene group. Violin plots indicate range of values, with a bar at median. P-value is calculated using a Mann-Whitney test. C) The 3′ end locations of oligoadenylated RNAs identified in exosome-depleted cells are shown at mRNA genes that are upregulated or unchanged by IntS9-depletion. (D) Kal1 (CG6173) locus displaying profiles of ChIP-seq for Integrator subunits, PRO-seq, and Start-seq following a time course of Triptolide treatment. (E) Decay rates for promoter Pol II were determined using Start-seq over a Triptolide treatment time course, and the percentage of upregulated or unchanged genes in each group is shown. (F-G) Average distribution of (F) H3K36me3 and (G) H3K4me3 ChIP-seq signal is shown, aligned around mRNA TSSs. Genes shown are those upregulated or unchanged in the PRO-seq assay upon IntS9-depletion. See also Figure S6.

We considered that Integrator-mediated RNA cleavage should occur on nascent RNA that has exited the polymerase. The structure of paused elongation complexes (Core and Adelman, 2019; Henriques et al., 2013; Vos et al., 2018), indicates that RNA emerges from the exit channel and is available for binding ∼15-20 nt upstream of the 3’ end position of the nascent RNA. Accordingly, the peak of oligoadenylated RNA 3’ end locations at upregulated genes is +35 nt (Figure 6C), which is 20nt upstream of the peak of paused Pol II at these genes, at +55nt (Figure S6A). From these data, we conclude that Integrator-repressed genes undergo markedly higher levels of Pol II termination as compared to non-Integrator target genes, and that promoter-proximally paused Pol II is the predominant target of Integrator-mediated RNA cleavage activity.

We next compared the stability of promoter-associated Pol II at Integrator-repressed genes after treatment with Triptolide. Based on increased premature termination at these genes, and our identification of Integrator enrichment at genes with unstable Pol II (Figure 1D), we predicted that Integrator-repressed genes would exhibit reduced promoter Pol II stability as compared to Integrator-unaffected genes. In agreement with this, we observed that Pol II was lost quickly at a majority of IntS9-repressed genes, with half-lives <10 minutes (Figures 6D and 6E). In contrast, genes whose expression is unchanged by IntS9-depletion presented a Pol II that is stable after Trp treatment, indicative of long-lived pausing (Figure 6E). Furthermore, genes upregulated by IntS9-depletion exhibited lower levels of H3K36me3 and H3K4me3 (Figures 6F, 6G and S6B) and higher levels of H3K4me1 (Figure S6C) than unchanged genes, consistent with defects in productive elongation. Thus, based on many independent lines of evidence we conclude that genes with unstable Pol II recruit Integrator, rendering them susceptible to promoter-proximal termination, and resulting in reduced productive RNA synthesis and chromatin features that accompany transcription elongation.

### Integrator-mediated gene repression is conserved in human cells

Our data in *Drosophila* indicate a mechanistically conserved role for Integrator in promoter-proximal termination of mRNA and eRNA synthesis. Although our model is in agreement with data from mammalian systems as regards eRNA biogenesis (Lai et al., 2015), it differs considerably from any of the proposed roles of Integrator at mammalian protein-coding genes (Barbieri et al., 2018; Gardini et al., 2014; Lai et al., 2015; Skaar et al., 2015; Stadelmayer et al., 2014). In particular, a majority of models posit that mammalian Integrator is an activator of transcription, and none of the proposed functions involve the IntS11 endonuclease in termination. For example, based on genomic studies of Integrator localization and activity in HeLa cells, it was proposed that Integrator stabilizes paused Pol II and facilitates both processive transcription elongation and RNA processing (Stadelmayer et al., 2014). Alternatively, other work in HeLa cells has implicated Integrator as critical for the rapid, EGF-mediated induction of ∼100 ‘immediate early’ genes, including *JUNB* and *FOS*. At these genes, Integrator was found to stimulate gene activity through recruitment of the Super Elongation Complex (Gardini et al., 2014). However, a detailed analysis of *JUNB* and several other immediate early genes gene in Integrator-depleted HeLa cells prior to EGF stimulation indicated that these genes were upregulated by loss of Integrator. Thus, it was suggested that Integrator inhibits expression of EGF-responsive genes under basal conditions (Skaar et al., 2015). Thus, it remains an open question whether, in the absence of a stimulus, mammalian Integrator plays a repressive role similar to that uncovered for the *Drosophila* complex.

To investigate whether loss of mammalian Integrator led to upregulation of gene transcription, as we observed for *Drosophila*, we analyzed previously published chromatin-associated RNA-seq from control and IntS11-depleted HeLa cells harvested prior to EGF stimulation. While chromatin-associated RNA-seq lacks the spatial resolution of PRO-seq, it is a significantly better indicator of ongoing transcription than is steady-state RNA-seq. Thus, we probed for differentially transcribed genes following IntS11-depletion in chromatin RNA-seq, using the same strategies employed for analysis of PRO-seq. Strikingly, we found a substantial number of genes upregulated in IntS11-depleted cells (N=667; Figures 7A and S7A), comparable to the number of genes downregulated under these conditions (N=616). Thus, mammalian Integrator appears capable of repressing as well as activating gene transcription. Importantly, despite the lower resolution of chromatin RNA-seq, increased transcript levels in Integrator-depleted cells are apparent within the initially transcribed region (Figure 7A), as observed in the *Drosophila* system.

**Figure 7.**
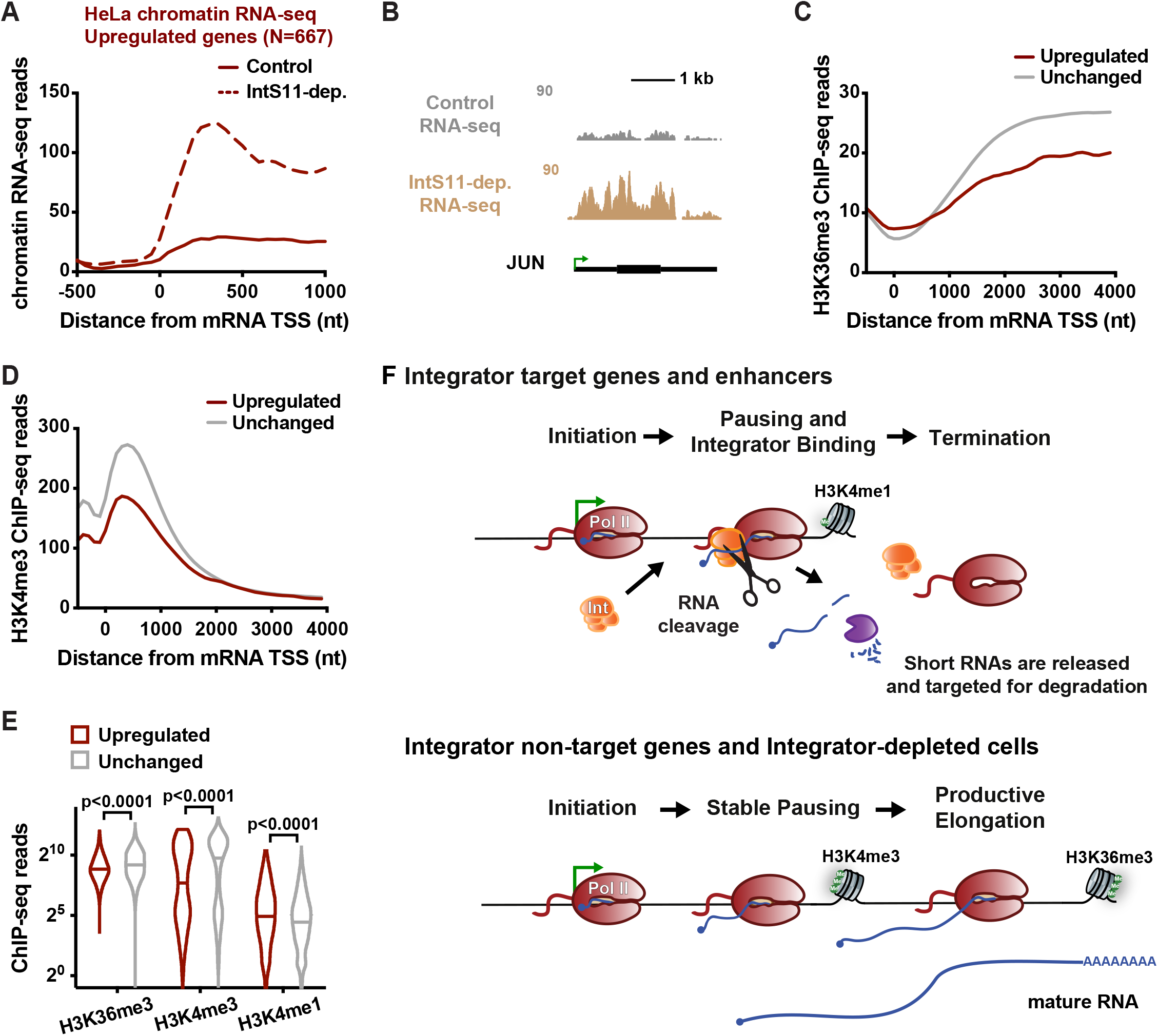
The Integrator complex represses expression of mammalian protein-coding genes. (A) Average distribution of chromatin RNA-seq reads in control and IntS11-depleted HeLa cells is shown for genes upregulated upon IntS11-depletion (data from Lai *et al*., 2015). (B) JUN locus showing upregulation of transcription upon IntS11-depletion. Shown are profiles of chromatin RNA-seq in control and IntS11-depleted HeLa cells (data from Lai *et al*., 2015). (C-D) Average distribution of (C) H3K36me3 and (D) H3K4me3 histone modifications (data from ENCODE project) is shown around mRNA TSSs for Upregulated (N=667) and unchanged (N=15979) genes. (E) H3K36me3, H3K4me3 and H3K4me1 ChIP-seq levels are shown for upregulated and unchanged genes. Violin plots show range of values, with a line indicating median. P-values are calculated using a Mann-Whitney test. (F) Schematic representation of the effect of the Integrator complex at protein-coding and enhancer loci. See also Figure S7.

The *JUNB* gene, which is a defined target of Integrator (Gardini et al., 2014), is strongly upregulated in HeLa cells depleted of IntS11 (Figure 7B), consistent with earlier work (Skaar et al., 2015). Moreover, many characterized immediate early genes exhibit elevated transcription under IntS11-depleted conditions and enriched Gene Ontology categories for upregulated transcripts include receptor and EGF pathways (Figure S7B). Interestingly, there is a concordance between upregulated pathways in *Drosophila* and human cells (compare Figure S7B to S4A), supporting a functional conservation of Integrator activity within specific pathways. Critically, these findings suggest that basal upregulation of stimulus-responsive genes upon Integrator depletion may be linked to the defective induction of these genes upon activation of signaling cascades.

To further probe the parallels between Integrator-mediated gene repression in *Drosophila* and human cells, we determined whether Integrator-repressed human genes also displayed chromatin features indicative of defective transcription elongation, such as reduced H3K36me3 and H3K4me3. As is seen in *Drosophila* (Figure 1B), both of these histone modifications were significantly lower at human genes upregulated upon Integrator depletion as compared to unchanged genes (Figures 7C and 7D). In addition, these genes showed enrichment in H3K4me1, a feature of both *Drosophila* Integrator gene targets and enhancers (Figures 7E and S7D). Thus, the significant commonalities among *Drosophila* and human genes repressed by Integrator, suggest a conserved mechanism across metazoan species (Figure 7F), wherein Integrator targets promoter-proximal elongation complexes at a set of genes to repress gene activity.

## DISCUSSION

Collectively, our results demonstrate that the Integrator complex mediates transcription attenuation in metazoan cells. This activity involves the association of Integrator with promoter-proximally paused Pol II, cleavage of nascent mRNA transcripts by the Integrator endonuclease, and promoter-proximal termination (Figure 7F). This inhibitory function is broad: 15% of *Drosophila* genes and enhancers are impacted by Integrator, with receptor, growth and proliferative pathways particularly affected. Furthermore, the mammalian Integrator complex targets genes in similar pathways for transcriptional repression, underlining the conserved nature of this behavior.

These data resolve long-standing questions about the intrinsic stability of promoter-proximal Pol II. We demonstrate that genes that harbor highly unstable promoter Pol II are those where there is an active process of termination, catalyzed by the Integrator complex. Our data support a model wherein the paused polymerase is inherently stable in the absence of termination factors, consistent with a wealth of biochemical characterization of elongation complexes (Kireeva et al., 2000; Wilson et al., 1999). Thus, we propose that rapid turnover of promoter Pol II at specific genes results from a regulated process of Integrator-mediated RNA cleavage and active dissociation of Pol II from the DNA template.

The mechanistic activity we uncover here for Integrator at protein-coding genes and enhancers parallels that described at snRNA genes, where Integrator cleaves the nascent RNA and promotes Pol II termination (Baillat and Wagner, 2015; Cazalla et al., 2011; Hernandez, 1985; Xie et al., 2015). Therefore, our model for Integrator function is parsimonious with its previously defined biochemical activities. Moreover, consistent with IntS9 and IntS11 subunits being paralogs of CPSF100 and CPSF73, respectively, there are many similarities between premature Pol II termination caused by Integrator, and mRNA cleavage and termination by the CPA machinery. We note that mRNA cleavage and termination at gene ends is coupled with polyadenylation to protect the released mRNA. Likewise, Integrator-catalyzed cleavage of snRNAs is coupled to proper 3’ end biogenesis. In contrast, termination driven by Integrator at protein-coding and enhancer loci would typically be followed by RNA degradation (Ogami et al., 2017). These results indicate that the Integrator endonuclease activity can be deployed for different purposes at different loci, with the outcome governed by the locus-specific recruitment of RNA processing or RNA decay machineries. Therefore, probing the interplay between Integrator and the complexes that govern RNA fate is an area that merits future study.

It has been established that cleavage and termination by the CPA machinery is greatly facilitated by pausing of Pol II (Proudfoot, 2016), as is snRNA 3’end formation by Integrator (Guiro and Murphy, 2017). Current models invoke a kinetic competition between Pol II elongation and termination, wherein slowed transcription elongation provides a greater window of opportunity for termination to occur (Fong et al., 2015; McDowell et al., 1994). Consistent with these models, we find that promoter-proximally paused Pol II is an optimal target for Integrator-mediated cleavage and termination at mRNA and eRNA loci. Our findings thus suggest a novel function for Pol II pausing in early elongation, wherein pausing provides a regulatory opportunity that enables gene attenuation.

It is interesting that Integrator-repressed genes, which exhibit very low levels of productive elongation, have chromatin characteristics that are common at enhancers. In particular, these genes display low levels of active histone modifications H3K4me3 and H3K36me3, with an enrichment in H3K4me1. Like at Integrator-repressed genes, transcription at enhancers is known to be non-productive, with a highly unstable Pol II that yields only short, rapidly degraded RNAs (Henriques et al., 2018; Kim and Shiekhattar, 2015). Thus, our data support models wherein these chromatin features reflect the level and productivity of transcription at the locus, rather than specifically demarcating the coding vs. non-coding potential of the region (Andersson et al., 2015; Core et al., 2014; Henriques et al., 2018; Soares et al., 2017).

Taken together, the role we describe here for Integrator in determining the fate of promoter Pol II sheds new light on Integrator function in development and disease states. Mutations in Integrator have been associated with a myriad of diseases (Rienzo and Casamassimi, 2016), with each of the 14 Integrator subunits implicated in one or more disorders. Intriguingly, many of these disease states are not characterized by defects in splicing and are often associated with disruption in normal development (Rienzo and Casamassimi, 2016). Thus, the human genetics foretold that Integrator functions extend well beyond snRNA processing. Accordingly, we find that Integrator targets a set of stimulus- and developmentally-responsive genes to potently repress their activity. It will be interesting in future work to tease out the specific roles of the individual Integrator subunits in gene regulation, in the hopes of exploiting this knowledge for therapeutic benefit.

## ACKNOWLEDGEMENTS

We thank Todd Albrecht for his help in generating *Drosophila* Integrator antibodies, Erik Andrulis for the Rrp40 antibody, William K. Russel and the UTMB Proteomics Core, and other members of the Adelman, Wilusz and Wagner labs for helpful discussions. Supported by National Institutes of Health grants R35-GM119735 (to J.E.W.), K99-GM131028 (to D.C.T.), Welch Foundation grant H-1889 (to E.J.W.); Startup Funds provided by Harvard Medical School (to K.A).

## AUTHOR CONTRIBUTIONS

T.H., N.D.E., K.L.H., K.A., and E.J.W. designed and performed RNA-seq, ChIP, ChIP-seq, and PRO-seq experiments; T.H., N.D.E., and K.A. analyzed genomic datasets; J.E.W. and D.C.T. performed Northern blots and provided insights. K.A. and E.J.W. wrote the manuscript with input from all authors.

## SUPPLEMENTAL FIGURE TITLES AND LEGENDS

**Figure S1. Related to Figure 1. mRNA TSSs occupied by unstable Pol II have enhancer-like histone modifications and are enriched in Integrator binding.** Active Drosophila genes were separated into groups based on the Promoter Pol II decay rate determined in Triptolide-treated cells (Henriques *et al*., 2018). All panels in this figure depict the same sets of genes, with consistent color-coding. Violin plots show range of values, with a line indicating median. P-values are calculated using a Mann-Whitney test.

(A) Violin plots showing Promoter (±150 nt from TSS) and Gene Body (+250 nt to +1250 nt from TSS) PRO-seq read counts, indicating similar levels of promoter Pol II, but less actively engaged Pol II at genes with the fastest Pol II decay rates.

(B) Average distributions (left) and levels (right; −200bp to +100bp from TSS) of ATAC-seq signal. n.s. means not significant.

(C) H3K36me3 (0bp to +500bp from TSS) ChIP-seq read counts are shown for genes with the fastest Pol II decay as compared to genes with more stable pausing of Pol II.

(D-E) Average distributions of H3K4me1 (left) and H3K4me3 (right) ChIP-seq signals, as metagene profiles and violin plots of signal (0bp to +500bp from TSS). Genes with rapid Pol II decay display higher H3K4me1/me3 ratios than other genes, such that many would be characterized as enhancers using standard ENCODE/ ChromHMM parameters.

(F) IntS1 (−250bp to +250bp from TSS) ChIP-seq signals at genes with the fastest Pol II decay versus other gene groups.

(G) Average distribution of IntS12 ChIP-seq signal, and violin plot of reads (−250bp to +250bp from TSS) at genes with the fastest Pol II decay versus those with more stable Pol II.

**Figure S2. Related to Figure 2. The Integrator complex attenuates expression of protein-coding genes.**

(A) Representative Western blot showing levels of IntS9 in cells following 60 h treatment with a control dsRNA, or a dsRNA targeting IntS9. Alpha-tubulin was used as a loading control.

(B) Representative Northern blots showing that mature snRNA levels are not affected upon IntS9-depletion. U6 snRNA biogenesis does not require the Integrator complex and serves as a loading control.

(C) RT-qPCR validation of representative genes affected by Integrator. mRNA levels from control and IntS9-depleted cells were normalized to RpS17 expression and the fold change upon IntS9 depletion is shown (mean ± SD, N=3).

(D-E) Gene Ontology analysis was performed on (D) the 723 transcripts upregulated or

(E) the 163 transcripts downregulated by IntS9-depletion in the RNA-seq assay. This corresponded to 511 and 111 unique genes, respectively, that had defined Gene Ontology ID annotations in DAVID Tools (v6.8). The top four enriched functional categories, pathways and domains are reported.

(F) Shown are promoter (±150 nt from TSS) and gene body (+250 to +1250 nt from TSS) PRO-seq signal density at genes upregulated by IntS9-depletion, as compared to unchanged genes. Normalized RNA-seq counts are also shown. Violin plots depict range of values, with line indicating median. P-values are from Mann-Whitney test.

(G) Overlap between IntS9-affected genes in RNA-seq and PRO-seq assays. P-values were calculated using a hypergeometric test using a total of 9,499 active mRNA genes.

**Figure S3. Related to Figure 3. IntS11 catalytic activity is essential for attenuation of protein-coding genes.**

(A) Representative Western blot showing protein levels of endogenous IntS11 and the exogenous IntS11 WT and E203Q transgenes. Cells were treated with a control dsRNA or a dsRNA targeting IntS11 for 60 h. Alpha-tubulin was used as a loading control.

(B) Representative Northern blots showing that mature snRNA levels are not affected upon IntS11-depletion. U6 snRNA biogenesis does not require the Integrator complex and serves as a loading control.

(C) Comparison of the fold changes in steady-state RNA-seq signals upon IntS9 or IntS11-depletion is shown. Pearson correlations were calculated separately for upregulated and downregulated genes, indicating good agreement between signals for upregulated genes.

(D) 2-D principal component analysis of RNA-seq read counts at mRNA genes. Note that the Control and IntS11-depleted cells rescued with WT IntS11 are grouped together in Principle component 1 (PC1), which describes the majority (82%) of the variance among samples.

(E) RT-qPCR validation of representative genes supporting mRNA attenuation and rescue of expression by IntS11. Stably transfected cells were treated for 60 h with a control dsRNA or dsRNA targeting IntS11. mRNA levels were normalized to RpS17 expression and the fold change is shown upon IntS11 depletion and the different rescue conditions (mean ± SD, N=3).

**Figure S4. Related to Figure 4. Loss of integrator complex causes transcription upregulation at protein-coding genes and enhancers.**

(A-B) Gene Ontology analysis of (A) the 1204 transcripts upregulated or (B) the 210 transcripts downregulated by IntS9-depletion, as defined from PRO-seq data. This corresponded to 1141 and 203 unique genes, respectively, that had defined Gene Ontology ID annotations in DAVID Tools (v6.8). The top four enriched functional categories, pathways and domains are reported. A single KEGG pathway was found to be enriched in downregulated genes.

(C) At left: Violin plots depict the change in promoter PRO-seq signal (± 150nt) upon IntS9-depletion for IntS9-affected genes (defined as in A) and unchanged genes (N=8085). Violin plots show range of values, with a line indicating median. At right: Difference in PRO-seq signal between IntS9-depleted and control cells is shown at downregulated (N=210) and unchanged genes (N=8085).

(D) Normalized PRO-seq signal at IntS9-upregulated enhancer RNA loci is shown downstream of enhancer TSSs. Upregulated eTSSs are defined from PRO-seq data from eTSS to +500bp, using P<0.05 and fold change >1.3.

(E) Difference in PRO-seq signal between IntS9-depleted and control cells is shown for enhancer loci that show no significant changes in PRO-seq signal upon IntS9-depletion (N=1232).

**Figure S5. Related to Figure 5. The Integrator complex is broadly associated with snRNA and mRNA genes.**

(A) Left: Signal from ChIP-seq for IntS1 versus IntS12 is shown at all 31 snRNA genes (using window from −150 to +350 bp from each TSS). Pearson correlation coefficient is indicated. Right: Average distribution of IntS12 and IntS1 ChIP-seq signal is depicted at snRNA loci. Read counts were summed in 25-nt bins, aligned on the TSS.

(B) Heatmap representations of IntS12 and IntS1 ChIP-seq signal and PRO-seq signal around mRNA TSSs (N=9499). Genes are ranked ordered by decreasing promoter-proximal IntS1 ChIP-seq signal (calculated from ±250 bp from TSS). Color bar at right indicates location of upregulated (red), downregulated (green) and unchanged (white) genes, based on PRO-seq in IntS9-depleted cells. Green arrow depicts TSS. Pearson correlation coefficients of promoter signals are shown below each heatmap.

(C) Distribution of IntS12 ChIP-seq signal along the transcription units of all active mRNA genes from TSS to transcription end site (TES), with bin size scaled by gene length.

(D) Average distribution of IntS12 ChIP-seq signal is shown at upregulated and unchanged genes.

(E) Average distribution of IntS1 and IntS12 ChIP-seq signal at IntS-bound eTSSs (N=691) vs. unbound (N=4182).

(F) Difference in PRO-seq signal between IntS9-depleted and control cells for IntS-bound and unbound eTSSs.

(G) ChIP-qPCR validation of IntS11 and IntS12 at promoters of representative Integrator target loci and snRNAs. ChIP signal is normalized relative to input (mean ± SD, N=3).

**Figure S6. Related to Figure 6. Integrator-repressed genes exhibit chromatin features consistent with unstable Pol II pausing and defective transcription elongation.**

(A) Average distribution of PRO-seq signal depicting the 3’ ends of nascent RNA held within elongation complexes (blue), as compared to the 3’ end locations of oligoadenylated RNAs identified in exosome-depleted cells (red). Shown are reads at genes upregulated by IntS9-depletion. The peak position of each distribution is indicated by an arrow.

(B) H3K36me3 (left) and H3K4me3 (right) ChIP-seq read counts are shown for genes upregulated or unchanged upon IntS9-depletion. Gene sets are defined as in Figure 4A. Windows used comprise TSS to +2000bp for H3K36me3, or TSS to +500bp from TSS for H3K4me3.

(C) At left: average distribution of H3K4me1 ChIP-seq signal is shown, aligned around TSSs, at upregulated or unchanged genes. At right: H3K4me1 ChIP-seq read counts (TSS to +500bp) are shown.

**Figure S7. Related to Figure 7. The Integrator complex affects expression of mammalian protein-coding genes under basal conditions.**

(A) Chromatin RNA-seq data from Lai *et al.,* 2015 were analyzed similarly to PRO-seq. Data are from HeLa cells depleted of IntS11 for 72h, as compared to cells expressing a scrambled shRNA. Depth normalized chromatin RNA-seq signal is shown. Significantly affected genes are defined using P<0.0001 and fold change >1.5.

(B-C) Gene Ontology analysis was performed on (B) the 667 genes upregulated or (C) the 616 genes downregulated by IntS11-depletion. This corresponded to 658 and 595 unique genes, respectively, that had defined Gene Ontology ID annotations in DAVID Tools (v6.8). The top four enriched functional categories, pathways and domains are reported.

(D) Average distribution of H3K4me1 ChIP-seq signal is shown, aligned around TSSs for upregulated or unchanged genes, defined as in A.

## SUPPLEMENTAL TABLE TITLES AND LEGENDS

**Table S1. Related to Figure 2. Integrator-affected genes identified by RNA-seq.**

List of genes that were identified as upregulated (N=723) or downregulated (N=163) in RNA-seq experiments performed on Control and IntS9-depleted cells. The threshold used to define IntS9-affected genes was p<0.0001 and fold change >1.5. Mean normalized counts in cells treated with control dsRNA or dsRNA targeting IntS9 is shown for each gene, along with the corresponding adjusted p-value.

**Table S2. Related to Figure 3. Integrator-affected genes identified by PRO-seq.**

List of genes that were identified as upregulated (N=1204) or downregulated (N=210) in the PRO-seq experiments performed on Control and IntS9-depleted cells. The threshold used to define IntS9-affected genes was p<0.0001 and fold change >1.5. Mean normalized counts in cells treated with control dsRNA or dsRNA targeting IntS9 is shown for each gene, along with the corresponding adjusted p-value.

**Table S3. Oligonucleotide sequences.** The oligonucleotide sequences used for RT-qPCR, ChIP-qPCR, plasmid cloning, Northerns and dsRNA synthesis are provided.

## METHODS

### Drosophila cell lines

*Drosophila* DL1 cells were cultured at 25°C in Schneider’s *Drosophila* medium (Thermo Fisher Scientific 21720024), supplemented with 10% (v/v) fetal bovine serum (HyClone SH30910.03), 1% (v/v) penicillin-streptomycin (Thermo Fisher Scientific 15140122), and 1% (v/v) L-glutamine (Thermo Fisher Scientific 35050061). Drosophila S2 cells from the DGRC were grown in Shields and Sang M3 (Sigma S3652) media supplemented with bactopeptone (BD Biosciences 211677), yeast extract (Sigma Y1000) and 10% FBS (Thermo Fisher Scientific 16000).

### Expression plasmid construction and generation of stable cell lines

To generate the selectable IntS11 expression plasmids, the previously described pUB-3xFLAG vector (Chen et al., 2012) Flag tag and MCS was cloned into the pMT-puro expression plasmid (a gift from David Sabatini, Addgene plasmid # 17923). *Drosophila* cDNA or the cDNA for the eGFP protein was then cloned into the resultant expression plasmid. The PCR primers are provided in Table S3. The IntS11 E203Q mutation (GAG to CAG) was subsequently introduced using site-directed mutagenesis. All plasmids were sequenced to confirm identity.

To generate DL1 cells stably maintaining the Flag-tagged IntS11WT, E203Q mutant, and the eGFP control line transgenes, 2 × 10^6^ cells were first plated in complete media in 6-well dishes. After 1 hour, 2 μg of pUB Flag-IntS11WT-puro, pUB Flag-IntS11E203Q-puro, or Flag-eGFP-puro were transfected using Fugene HD (Promega E2311). On the following day, 2.5 μg/mL puromycin was added to the media to select and maintain the cell population.

### RNAi

Double-stranded RNAs from the DRSC (*Drosophila* RNAi Screening Center) were generated by *in vitro* transcription (MEGAscript kit, Thermo Fisher Scientific AMB13345) of PCR templates containing the T7 promoter sequence on both ends. Primer sequences are provided in Table S3. Knockdown experiments in 6-well dishes were then performed by bathing 1.5×10^6^ cells with 2 μg of dsRNA, followed by incubation for 60 hours of standard cell culture conditions. For RNAi + rescue experiments (Figure 3) cells were incubated for 60 hours in the presence of dsRNA and media was supplemented with a final concentration of 100 μM CuSO4 to induce expression of the RNAi-resistant IntS11 WT or IntS11 E203Q transgenes.

### RT-qPCR

Total RNA was isolated using Trizol and cDNA was reverse transcribed using M-MLV Reverse Transcriptase (Thermo Fisher Scientific 28025) according to the manufacturer’s instructions. Random hexamers were used for cDNA synthesis and RT-qPCR was then carried out in triplicate using Bio-Rad iTaq Universal SYBR Green Supermix (Bio-Rad 1725120). All RT-qPCR primers are provided in Table S3.

### Analysis of protein expression by Western blotting and immunofluorescence

For Western blotting, cells were gently washed in PBS and then resuspended in RIPA buffer (150 mM NaCl, 1% Triton X-100, 50 mM Tris pH 7.5, 0.1% SDS, 0.5% sodium-deoxycholate, and protease inhibitors [Roche 11836170001]). Lysates were passed 10 times through a 28.5 gauge needle and cleared by centrifugation at 20,000x*g* for 20 min at 4°C. Lysates were then resolved on a NuPAGE 4-12% Bis-Tris gel (Thermo Fisher Scientific NP0323) and transferred to a PVDF membrane (Bio-Rad 1620177). Primary antibody incubations (IntS9 [guinea pig], IntS11 [rabbit] (Ezzeddine et al., 2011) or alpha-tubulin (rabbit, abcam ab15246) were all done at room temperature for 2 hours with a 1:1000 dilution in 5% milk in TBS-0.1% Tween. Conjugated secondary antibodies against rabbit (GE Healthcare NA934) or guinea pig (Sigma AP108P) were incubated at room temperature for 90 minutes with 1:10000 dilution in TBS-0.1% Tween. Membranes were processed using SuperSignal West Pico Chemiluminescent Substrate (Thermo Fisher Scientific PI34080).

### Northern blotting

Total RNA was isolated using Trizol (Thermo Fisher Scientific 15596018) as per the manufacturer’s instructions. Small RNAs were separated by 8% denaturing polyacrylamide gel electrophoresis (National Diagnostics EC-833) and electroblotted/UV crosslinked to Hybond N+ membrane (GE Healthcare RPN303B). ULTRAhyb-oligo hybridization Buffer (Thermo Fisher Scientific AM8663) was used as per the manufacturer’s instructions. All oligonucleotide probe sequences are provided in Table S3. Blots were viewed and quantified with the Typhoon 9500 scanner (GE Healthcare) and quantified using ImageQuant (GE Healthcare). Representative blots from ≥3 experiments are shown.

### Chromatin Immunoprecipitation (ChIP)-qPCR

A 10-cm dish of 5 x 10^7^ DL1 cells was harvested into a 15 mL tube and centrifuged at 1,500x *g* for 2 min. Cells were then washed with 10 mL PBS and centrifuged at 1,500x *g* for 2 min. The cell pellet was resuspended in 10 mL of Fixing Buffer (50 mM Hepes pH 7.5, 100 mM NaCl, 1 mM EDTA pH 8.0, 0.5 mM EGTA pH 8.0 with 1% formaldehyde) and incubated at room temperature for 30 min. 0.5 mL of 2.5 M glycine was then added (final concentration of 0.125 M) and incubated at room temperature with rotation for 5 min, centrifuged at 1,500 *g* for 2 min, and washed two times with 10 mL PBS. Cells were lysed using lysis buffer (50 mM HEPES pH 7.9, 140 mM NaCl, 1 mM EDTA, 10% glycerol, 0.5% NP-40, 0.25% Triton X-100) for 10 min on ice and centrifuged at 1,500 *g* for 2 min. The pellet was then washed 2x in Wash Buffer (10 mM Tris-HCl pH 8.1, 200 mM NaCl, 1 mM EDTA pH 8.0, 0.5 mM EGTA pH 8.0) and resuspended in 1 mL Shearing Buffer (0.1% SDS, 1 mM EDTA, 10 mM Tris-HCl pH 8.1). The suspension was sonicated at 4°C using a Covaris S220 machine to obtain 500 bp DNA fragments in TC12×12 tubes with AFA fiber (Settings: Time-15 min, Duty Cycle-5%, Intensity-4, Cycles per Burst-200, Power mode Frequency-Sweeping, Degassing mode-Continuous, AFA Intensifier-none, Water level-8). To the 1 mL of sheared chromatin, 115 μL of 10% Triton X-100 and 34 μL 5 M NaCl was added per ml of sheared chromatin, so that the final concentration of the sample is 1% Triton X-100 and 150 mM NaCl. Sheared chromatin was pre-cleared with protein A/G beads and 10 μL was reserved as input control. For each IP sample, 100 μL of sheared chromatin was diluted to 1 mL using IP Buffer (0.1% SDS, 1 mM EDTA, 10 mM Tris-HCl pH 8.1, 1% Triton X-100, 150 mM NaCl) and incubated overnight at 4°C with 10 μL of serum. The next day, lysates were immunoprecipitated with protein A/G beads for 2 h at 4°C and washed once with low salt buffer (0.1% SDS, 1% Triton X-100, 2 mM EDTA, 20 mM Hepes pH 7.9, 150 mM NaCl), twice with high salt buffer (0.1% SDS, 1% Triton X-100, 2 mM EDTA, 20 mM Hepes pH 7.9, 500 mM NaCl), once with LiCl buffer (100 mM Tris-HCl pH 7.5. 0.5 M LiCl, 1% NP-40, 1% Sodium Deoxycholate), and once with TE. Immunocomplexes were eluted and de-crosslinked at 65°C overnight with Proteinase K and RNase A. DNA was extracted by phenol-chloroform and ethanol precipitated. DNA was resuspended in 100 uL, and 2 uL was used for each qPCR reaction.

### Quantification and Statistical Analysis

For RT-qPCRs statistical significance for comparisons of means was assessed by Student’s t test. Unless otherwise indicated, the comparison was to the control RNAi treated samples. Statistical details and error bars are defined in each figure legend.

### Genomic Data Availability

All datasets generated in this study are available for download from GEO (GSE114467). Start-seq from untreated S2 cells and Start-seq from Rrp40-depleted S2 cells was published previously (Henriques et al., 2013) and is available for download from GEO (GSE49078). Start-seq from Triptolide-treated S2 cells was published previously (Krebs et al., 2017) and is available for download from GEO (GSE77369). H3K36me3, H3K4me3 and H3K4me1 ChIP-seq datasets from S2 cells were published previously (Henriques et al., 2018) and are available for download from GEO (GSE85191). Chromatin RNA-seq from Control and IntS11-dep. in HeLa cells was published previously (Lai et al., 2015) and is available for download from GEO (GSE68401). H3K36me3, H3K4me3 and H3K4me1 ChIP-seq datasets from HeLa-S3 cells are available as part of the ENCODE project (Gerstein et al., 2012) and can be retrieved under the following accession numbers GSM733711, GSM733682, GSM798322 from GEO (GSE29611).

### Generation of Transcript Annotations

All transcript annotations for *D. melanogaster* r5.57 were downloaded from flybase.org in GTF format and filtered such that only “exon” entries for the feature types considered for re-annotation remained. Annotations from chrY, chrM, and random chromosomes were also excluded. Unique “gene_id” values were assigned to each transcript, such that those grouped and represented by a single member in TSS-based analyses were identical. Precise TSS locations employed were based on high-resolution Start-seq data as described previously (Henriques et al., 2013; 2018; Nechaev et al., 2010). The start location of each transcript was adjusted to the observed TSS from Start-seq when this resulted in truncation, rather than extension of the model. If the observed TSS fell within an intron, all preceding exons were removed, and the transcript start was set to the beginning of the following downstream exon. Gene annotations for the human genome (hg19, GRCh37 genome build July 2019) were downloaded from gencodegenes.org in GTF format and filtered such that only “gene” entries for the “protein_coding” feature type remained. Annotations from chrM, and random chromosomes were also excluded.

### TSS clustering based on promoter Pol II half-lives upon Trp treatment

TSS clustering was accomplished as described in (Henriques et al., 2018) using k-medoids clustering based on the Clustering Large Applications (CLARA) object in R.

### Features associated with genes with short-lived promoter Pol II occupancy

A comprehensive repertoire of ChIP-seq datasets from (Baumann and Gilmour, 2017; Henriques et al., 2018; Kaye et al., 2018; Lim et al., 2013; Weber et al., 2014) and ChIP-chip from the modENCODE database (Ho et al., 2014; modENCODE Consortium et al., 2010) was used representing a total of 111 datasets that include transcription factors, chromatin remodelers and histone modifications.

To find features enriched at protein-coding transcription start sites with short-lived promoter Pol II occupancy a similar approach to the web-based tool ORIO (Lavender et al., 2017) was taken. Analysis of all datasets was anchored on the TSS locations of protein-coding transcripts based on high-resolution Start-seq data (see generation of transcript annotations above). A total of 8389 protein-coding TSSs, in which a decay rate could be calculated, was used. A rank order was given to the TSS feature list based on the decay rate clustering. Read coverage for each dataset used was determined at each TSS using a window that originates 500 nucleotides upstream of the TSS and extends downstream by twenty 50 nt non-overlapping bins, with total window size of 1000 nucleotides. Correlative analysis was then performed considering read coverage values. A total read coverage value was found for each genomic feature by adding the coverage from the datasets across all bins in a genomic window. Clustering methods were then applied to total read coverage values considering both the datasets and individual genomic features. To group datasets, the Pearson and Spearman correlation value for each pair of datasets was determined by comparing feature coverage values. To group the datasets, the correlation value for each pair of datasets is found by comparing feature coverage values. Datasets were then grouped by hierarchical clustering.

### ATAC-seq library generation and mapping

ATAC-seq libraries from 3 independent biological replicates were generated. 50,000 *Drosophila* S2 cells were incubated in CSK buffer (10 mM PIPES pH 6.8, 100 mM NaCl, 300 mM sucrose, 3 mM MgCl2, 0.1% Triton X-100) on ice for 5 min. An aliquot of 2.5 µl of Tn5 Transposase was added to a total 25 µl reaction mixture and genomic DNA was purified using a Qiagen MinElute PCR purification kit (Qiagen) following manufacturer’s instructions. After PCR amplification, DNA fragments were purified with AMPure XP (1:3 ratio of sample to beads). Libraries were sequenced using a paired-end 150 bp cycle run on an Illumina NextSeq 500

Paired-end reads were filtered for adapter sequence and low quality 3’ ends using cutadapt 1.14, discarding those containing reads shorter than 20 nt (-m 20 -q 10), and removing a single nucleotide from the 3’ end of all trimmed reads to allow successful alignment with bowtie 1.2.2 to the dm3 genome assembly. The parameters used in each alignment were: up to 2 mismatches, a maximum fragment length of 1000 nt, and uniquely mappable, and unmappable pairs routed to separate output files (-m1, -v2, -X1000, --un). Non-duplicate reads mapping uniquely to dm3, representative of short fragments (> 20 nt and < 150 nt), were separated, and fragment centers determined in 25 nucleotide windows resolution, genome-wide, and expressed in bedGraph format. Combined bedGraphs for all replicates were generated by summing counts per bin for all replicates.

| **Sample** | **Total reads** | **Uniquely mapped reads (Percentage of total)** | **Agreement between replicates (Spearman's rho)** |
| --- | --- | --- | --- |
| ATAC-seq | 42,095,224 | 62.94% | >0.97 |

### RNA-seq library generation and mapping

DL1 cells were treated for 60 h with a control (Beta-galactosidase) dsRNA or a dsRNA to deplete either IntS9 or IntS11 (see RNAi details above) followed by total RNA isolation with Trizol (Thermo Fisher Scientific 15596026) following manufacturer’s instructions. RNA quality was confirmed with a BioAnalyzer (Agilent). Using Oligo d(T)25 Magnetic Beads (NEB S1419S), polyA+ RNA from 2.5 μg of total RNA was then enriched and RNA-seq libraries prepared using the Click-seq library preparation method using a 1:35 azido-nucleotide ratio (Jaworski and Routh, 2018). Libraries were sequenced using a single-end 75 bp cycle run on an Illumina NextSeq 500.

Sequencing reads were filtered (requiring a mean quality score ≥20), trimmed to 50 nt, and then mapped to the dm3 reference genome using STAR 2.5.2b. Default parameters were used except that multimappers were reported randomly (outMultimapperOrder Random), spurious junctions were filtered (outFilterType BySJout), minimum overhang for non-annotated junctions was set to 8 nucleotides (alignSJoverhangMin 8), and non-canonical alignments were removed (outFilterIntronMotifs RemoveNoncanonicalUnannotated). The total number of RNA-seq reads aligned in the control, IntS9 or IntS11 RNAi samples is described in the table below.

| **Sample** | **Total Reads** | **Mappable Fragments** | **Agreement between replicates (Spearman's rho)** |
| --- | --- | --- | --- |

|  |  | (Percentage of total) |  |
| --- | --- | --- | --- |
| Control (βgal) | 65,094,896 | 71.58% | >0.98 |
| IntS9-dep. | 54,967,569 | 68.20% | >0.98 |
| Control (βgal) + eGFP | 69,413,816 | 87.42% | >0.99 |
| IntS11-dep. + eGFP | 49,878,632 | 84.54% | >0.99 |
| IntS11-dep. + WT | 51,586,717 | 85.29% | >0.99 |
| IntS11-dep. + E203Q | 57,653,391 | 86.64% | >0.99 |

### MISO Analysis

Mixture of Isoform analysis (MISO) (Katz et al., 2010) was performed using the latest stable build (ver. 0.5.4) following the directions for an exon-centric analysis on the documents section of the developer’s site (http://miso.readthedocs.io/en/fastmiso/). Differential expression was compared between the control (Beta-galactosidase) and IntS9-depleted RNA-seq BAM files for retained introns, skipped exons, alternative 5’ splice sites, alternative 3’ splice sites, and mutually excluded exons using the *Drosophila* annotations mentioned above. The results were then filtered using the developer suggested default settings to contain only events with: (a) at least 10 inclusion reads, (b) 10 exclusion reads, such that (c) the sum of inclusion and exclusion reads is at least 30, and (d) the ΔΨ is at least 0.25 with a (e) Bayes factor of at least 20, and (a)-(e) are true in one of the samples. Using this filter, locations of alternative splicing events were compared to Flybase annotated chromosomal regions using the UCSC genome browser table browser to identify the FBgnIDs of affected genes. The number of changes in splicing events are described in the table below.

| Splicing Event Type | Events compared | Events passing filter | Percent Events Passing Filter |
| --- | --- | --- | --- |
| Retained Intron | 24353 | 412 | 1.69% |
| Alternative 5'SS | 3231 | 63 | 1.95% |
| Alternative 3'SS | 1584 | 46 | 2.9% |
| Skipped Exon | 1376 | 27 | 1.96% |
| Mutually Exclusive Exon | 73 | 0 | 0% |

All Flybase genes that included any splicing event that passed filter in MISO were removed from the list of active genes, such that a total of 9,499 active genes were investigated for the effects of IntS9 depletion.

### Differentially expressed genes in RNA-seq

Read counts were calculated per gene, in a strand-specific manner, based on annotations described in the modified transcript annotations section above, using featureCounts (Liao et al., 2014). Differentially expressed genes were identified using DESeq2 v1.18.1(Anders and Huber, 2010) under R 3.3.1. For Control versus IntS9-depletion comparisons RNA-seq size factors were determined based on DESeq2 (Control [βgal]: 1.1861939, 1.4205182, 1.2440253; IntS9-dep.: 1.0780809, 0.9979663, 0.8519904), and at an adjusted p-value threshold of <0.0001 and fold-change > 1.5, 886 genes (out of 9499) were identified as differentially expressed upon IntS9 depletion in DL1 cells. For Control versus IntS11-depletion or rescue samples comparisons RNA-seq size factors were determined based on DESeq2 (Control [βgal]: 1.3346867, 1.8951248, 0.6622473; IntS11-dep.: 0.8673446, 0.9127478, 0.9793937; IntS11-dep. + WT rescue: 1.1305191, 1.0792675, 0.7458915; IntS11-dep. + E203Q rescue: 1.1589313, 1.1588886, 0.7106579) and fold-changes calculated. For Control versus IntS11-depletion chromatin RNA-seq size factors were determined based on DESeq2 (Control: 1.1315534, 1.1665893; IntS11-dep.: 0.8940834, 0.8515502;) and at an adjusted p-value threshold of <0.0001 and fold-change > 1.5, 1283 genes (out of 17262) were identified as differentially expressed upon IntS11 depletion in HeLa cells. UCSC Genome Browser tracks displaying mean read coverage were generated from the combined replicates per condition, normalized as in the differential expression analysis.

### Sequencing, mapping, and data analysis of ChIP-seq

For IntS1 and IntS12 ChIP-seq, DL1 cells were crosslinked for 30 min with 1% formaldehyde. Material was then sheared using the Covaris S220 system and immunoprecipitations for 3 (IntS1 and IntS12) independent biological replicates were carried out with 10 μl anti-IntS1 or anti-IntS12 antibodies per 3 x 10^7^ cells. Additionally, 3 independent biological replicates of input material were carried through. Immunoprecipitated and input material was phenol-chloroform purified and ChIP-seq libraries were prepared using the NEBNext Ultra II DNA library kit (NEB) according to the manufacturer’s instructions with 35ng of DNA of each sample. IntS1, IntS12 and input ChIP-seq libraries were then sequenced using a paired-end 75 bp cycle run on the Illumina NextSeq system with standard sequencing protocols. Raw sequences were aligned at full length against the dm3 version of the *Drosophila* genome using Bowtie version 1.2.2 (Langmead et al., 2009) with a maximum allowed mismatch of 2 (-m1 – v2). The yield of uniquely mappable reads for each set of biological replicates is listed below.

| **Sample** | **Total reads** | **Uniquely mapped reads (Percentage of total)** | **Agreement between replicates (Spearman's rho)** |
| --- | --- | --- | --- |
| Input | 76,003,314 | 62.82% | >0.97 |
| IntS1 | 92,977,225 | 61.76% | >0.97 |
| IntS12 | 107,536,665 | 62.66% | >0.97 |

Datasets were mapped as described above against the dm3 version of the *Drosophila* genome. The genomic location of mapped reads was compiled using custom scripts and visually examined using the UCSC genome browser in bedGraph format. ChIP-seq hit locations were filtered based on fragment length. The 3 biological replicates of each ChIP-seq dataset were combined and binned in 25 bp windows for visualization in bedGraph files. IntS1 and IntS12 were downsampled by a factor of 1.202985486 and 1.411913925, respectively to match the number of reads in the input dataset. To remove background signal, input signal was subtracted from IntS1 and IntS12 datasets and bedGraphs were generated with 25 bp windows for visualization.

### IntS1 and IntS12 ChIP-Seq peak calling and annotation

IntS1 and IntS12 ChIP-seq peaks were called with Homer (v4.9) using (-style factor) and input as background (-i). Filtering based on local signal was set to 3 (-L 3) and fold-change signal over input was also set to 3 (-F 3). 490 IntS1 and 553 IntS12 peaks were identified. A peak was assigned to enhancer TSSs (eTSSs) if the peak center would be within ± 500 bp from the eTSS. A total of 691 eTSSs were found to be bound by at least one Integrator subunit.

### Metagene analysis

Composite metagene distributions were generated by summing sequencing reads at each indicated position with respect to the TSS and dividing by the number of TSSs included within each group. These were plotted across a range of distances. Heatmaps were generated using Partek Genomics Suite version 6.15.0127.

### Identification of Start-seq reads with non-templated 3’ end residues

Start-seq from Rrp40-depleted S2 cells was published previously (Henriques et al., 2013) and is available for download from GEO (GSE49078). Data were analyzed as described previously (Henriques et al., 2013). Briefly, Start-RNA reads were trimmed to 26 nt and aligned to the *D. melanogaster* reference genome index with Bowtie version 1.2.2, maintaining unique alignments and allowing 2 mismatches (-m1 -v2). To account for the different depths of sequencing across the data sets, all data sets were normalized by uniquely mappable reads. To then identify Start-RNAs with non-templated 3’ end residues, reads that initially failed to align with the above Bowtie parameters were specifically trimmed at the 3’ end to remove terminal A nucleotides. Reads trimmed of at least 3 A’s with at least 18 nt remaining after trimming were aligned to the genome (note that reads with >26 nt remaining after trimming were further trimmed at the 5’ end to 26mers) and counted as uniquely-aligned Start-RNAs. The percentage and location of Start-seq reads ending in 3 or more A residues (out of total Start-seq reads mapping to that gene) was calculated for each gene in all the groups.

### PRO-seq library preparation and data analysis

DL1 cells treated for 60 h with a control (Beta-galactosidase) dsRNA or a dsRNA targeting IntS9 were permeabilized as described below. All temperatures were at 4°C or ice cold unless otherwise specified. Cells were washed once in ice-cold 1x PBS and resuspended in Buffer W (10 mM Tris-HCl pH 8.0, 10% glycerol, 250 mM sucrose, 10 mM KCl, 5 mM MgCl_2_, 0.5 mM DTT, protease inhibitors cocktail (Roche), and 4 u/mL RNase inhibitor [SUPERaseIN, Ambion]) at the cell density of 2 × 10^7^ cells/mL. 9x volume of Buffer P (10 mM Tris-HCl pH 8.0, 10% glycerol, 250 mM sucrose, 10 mM KCl, 5 mM MgCl2, 0.5 mM DTT, 0.1% Igepal, protease inhibitors cocktail (Roche), 4 u/mL RNase inhibitor [SUPERaseIN, Ambion]) was then immediately added. Cells were gently resuspended and incubated for up to 2 min on ice. Cells were then recovered by centrifugation (800 x *g* for 4 min) and washed in Buffer F (50 mM Tris-HCl pH 8.0, 40% glycerol, 5 mM MgCl_2_, 0.5 mM DTT, 4 u/mL RNase inhibitor [SUPERaseIN, Ambion]). Washed permeabilized cells were finally resuspended in Buffer F at a density of 1×10^6^ cells/30 μL and immediately frozen in liquid nitrogen. Permeabilized cells were stored in −80°C until usage.

PRO-seq run-on reactions were carried out as follows: 1 × 10^6^ permeabilized cells spiked with 5 × 10^4^ permeabilized mouse embryonic stem cells were added to the same volume of 2x Nuclear Run-On reaction mixture (10 mM Tris-HCl pH 8.0, 300 mM KCl, 1% Sarkosyl, 5 mM MgCl2, 1 mM DTT, 200 μM biotin-11-A/C/G/UTP (Perkin-Elmer), 0.8 u/μL SUPERaseIN inhibitor [Ambion]) and incubated for 5 min at 30°C. Nascent RNA was extracted using a Total RNA Purification Kit following the manufacturer’s instructions (Norgen Biotek Corp.). Extracted nascent RNA was fragmented by base hydrolysis in 0.25 N NaOH on ice for 10 min and neutralized by adding 1x volume of 1 M Tris-HCl pH 6.8. Fragmented nascent RNA was bound to 30 μL of Streptavidin M-280 magnetic beads (Thermo Fisher Scientific) in Binding Buffer (300 mM NaCl, 10 mM Tris-HCl pH 7.4, 0.1% Triton X-100). The beads were washed twice in High salt buffer (2 M NaCl, 50 mM Tris-HCl pH 7.4, 0.5% Triton X-100), twice in Binding buffer, and twice in Low salt buffer (5 mM Tris-HCl pH 7.4, 0.1% Triton X-100). Bound RNA was extracted from the beads using Trizol (Invitrogen) followed by ethanol precipitation.

For the first ligation reaction, fragmented nascent RNA was dissolved in H_2_O and incubated with 10 pmol of reverse 3’ RNA adaptor (5’p-rNrNrNrNrNrNrGrArUrCrGrUrCrGrGrArCrUrGrUrArGrArArCrUrCrUrGrArArC-/3’InvdT/) and T4 RNA ligase I (NEB) under manufacturer’s conditions for 2 h at 20°C. Ligated RNA was enriched with biotin-labeled products by another round of Streptavidin bead binding and washing (two washes each of High, Binding and Low salt buffers and one wash of 1x Thermo Pol Buffer (NEB)). To decap 5’ ends, the RNA products were treated with RNA 5’ Pyrophosphohydrolase (RppH, NEB) at 37°C for 30 min followed by one wash of High, Low and T4 PNK Buffer. To repair 5’ ends, the RNA products were treated with Polynucleotide Kinase (PNK, NEB) at 37°C for 30 min.

5’ repaired RNA was ligated to reverse 5’ RNA adaptor (5’-rCrCrUrUrGrGrCrArCrCrCrGrArGrArArUrUrCrCrA-3’) with T4 RNA ligase I (NEB) under manufacturer’s conditions for 2 h at 20°C. Adaptor ligated nascent RNA was enriched with biotin-labeled products by another round of Streptavidin bead binding and washing (two washes each of High, Binding and Low salt buffers and one wash of 1x SuperScript IV Buffer [Thermo Fisher Scientific]), and reverse transcribed using 25 pmol RT primer (5’-AATGATACGGCGACCACCGAGATCTACACGTTCAGAGTTCTACAGTCCGA-3’) for TRU-seq barcodes (RP1 primer, Illumina). A portion of the RT product was removed and used for trial amplifications to determine the optimal number of PCR cycles. For the final amplification, 12.5 pmol of RPI-index primers (for TRU-seq barcodes, Illumina) was added to the RT product with Phusion polymerase (NEB) under standard PCR conditions. Excess RT primer served as one primer of the pair used for the PCR. The product was amplified 12∼14 cycles and beads size selected (ProNex Purification System, Promega) before being sequenced in NextSeq 500 machines in a mid-output 150 bp cycle run.

PRO-seq libraries from 3 independent biological replicates (DL1 control (bgal) RNAi or IntS9 RNAi) were generated. Paired-end reads were trimmed to 42 nt, for adapter sequence and low quality 3’ ends using cutadapt 1.14, discarding those containing reads shorter than 20 nt (-m 20 -q 10), and removing a single nucleotide from the 3’ end of all trimmed reads to allow successful alignment with Bowtie 1.2.2. Remaining pairs were paired-end aligned to the mm10 genome index to determine spike-normalization ratios based on uniquely mapped reads. Mappable pairs were excluded from further analysis, and unmapped pairs were aligned to the dm3 genome assembly. Identical parameters were utilized in each alignment described above: up to 2 mismatches, maximum fragment length of 1000 nt, and uniquely mappable, and unmappable pairs routed to separate output files (-m1, -v2, -X1000, --un). Pairs mapping uniquely to dm3, representing biotin-labeled RNA 3’ ends, were separated, and strand-specific counts of the 3’ mapping positions determined at single nucleotide resolution, genome-wide, and expressed in bedGraph format with “plus” and “minus” strand labels swapped for each 3’ bedGraph, to correct for the “forward/reverse” nature of Illumina paired-end sequencing (see (Mahat et al., 2016)). Counts of pairs mapping uniquely to spike-in RNAs (mouse genome) were determined for each sample. Uniquely mappable reads were determined, and a normalization factor calculated. In this case, the samples displayed highly comparable recovery of spike-in reads, thus only normalization based on the DESeq2 size factors (see below) was used for each bedGraph. Combined bedGraphs were generated by summing counts per nucleotide of both replicates for each condition.

| **Sample** | **Total reads** | **Uniquely mapped reads (Percentage of total)** | **Agreement between replicates (Spearman's rho)** |
| --- | --- | --- | --- |
| Control (βgal) | 60,860,471 | 48.16% | >0.98 |
| IntS9-dep. | 57,112,558 | 53.29% | >0.99 |

Read counts were calculated per gene, in a strand-specific manner, based on annotations described in the modified transcript annotations section above, using featureCounts (Liao et al., 2014). This quantification procedure includes signal only in the gene body (+250 from TSS to annotated gene end). Differentially expressed genes were identified using DESeq2 v1.18.1 (Anders and Huber, 2010) under R 3.3.1. PRO-seq size factors were determined based on DESeq2 (for Control: 1.0029079, 1.2830936, 0.8962051; IntS9-dep.: 0.9151691, 0.9156818, 1.0672821). At an adjusted p-value threshold of <0.0001 and fold-change >1.5, 1,414 mRNA genes were identified as differentially expressed upon IntS9-depletion in DL1 cells. UCSC Genome Browser tracks displaying mean read coverage were generated from the combined replicates per condition, normalized as in the differential expression analysis.

### Genomic statistical tests

For RNA-seq, PRO-seq, and ChIP-seq experiments, statistical significance for comparisons was assessed by Mann-Whitney (pairwise tests) test. Statistical details and error bars are defined in each figure legend. To test for the significant overlap between IntS9-upregulated or IntS9-downregulated genes in RNA-seq and PRO-seq, a hypergeometric test was used from a total of 9499 active mRNA genes.

### Gene Ontology Analysis

Gene Ontology analysis was performed using DAVID (v6.8) online tool with standard parameters (https://david.ncifcrf.gov/home.jsp). The number of affected genes used to identify the top Biological categories and Pathways is described in the table below.

| **Treatment** | **Assay** | **Number Upregulated transcripts** | **Number Downregulated transcripts** |
| --- | --- | --- | --- |
| Control vs. IntS9-dep. | RNA-seq | 723 | 163 |
| Control vs. IntS9-dep. | PRO-seq | 1204 | 210 |
| Control vs. IntS11-dep. | Chromatin RNA-seq | 667 | 616 |

