## Supplemental Figures for "The Integrator complex terminates promoter-proximal transcription at protein-coding genes"

Figure S1

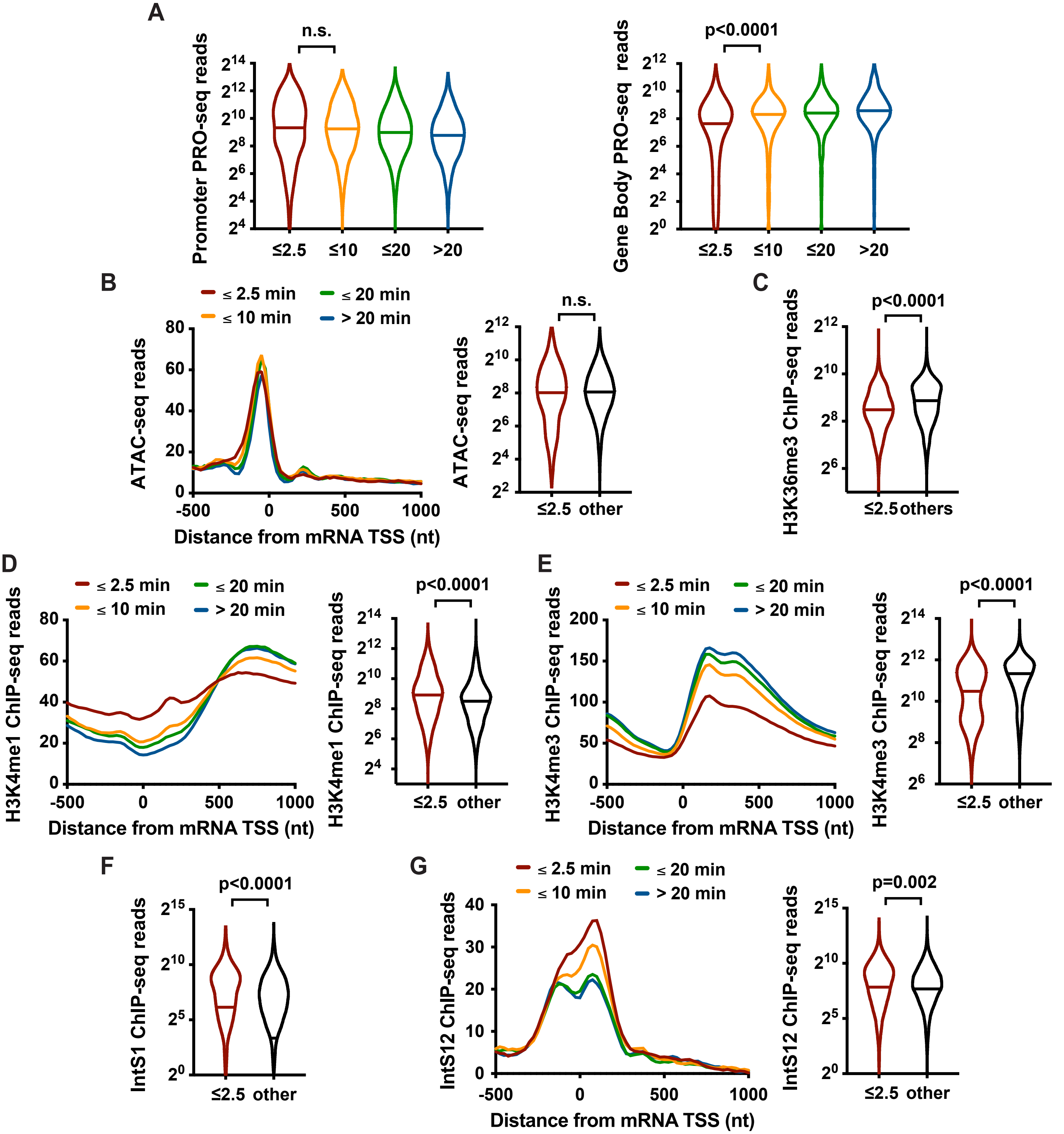


**Figure S1. Related to Figure 1. mRNA TSSs occupied by unstable Pol II have enhancer-like histone modifications and are enriched in Integrator binding.** Active Drosophila genes were separated into groups based on the Promoter Pol II decay rate determined in Triptolide-treated cells (Henriques *et al.,* 2018). All panels in this figure depict the same sets of genes, with consistent color-coding. Violin plots show range of values, with a line indicating median. P-values are calculated using a Mann-Whitney test.

(A) Violin plots showing Promoter (±150 nt from TSS) and Gene Body (+250 nt to +1250 nt from TSS) PRO-seq read counts, indicating similar levels of promoter Pol II, but less actively engaged Pol II at genes with the fastest Pol II decay rates.

(B) Average distributions (left) and levels (right; -200bp to +100bp from TSS) of ATAC-seq signal. n.s. means not significant.

(C) H3K36me3 (0bp to +500bp from TSS) ChIP-seq read counts are shown for genes with the fastest Pol II decay as compared to genes with more stable pausing of Pol II.

(D-E) Average distributions of H3K4me1 (left) and H3K4me3 (right) ChIP-seq signals, as metagene profiles and violin plots of signal (0bp to +500bp from TSS). Genes with rapid Pol II decay display higher H3K4me1/me3 ratios than other genes, such that many would be characterized as enhancers using standard ENCODE/ ChromHMM parameters.

(F) IntS1 (-250bp to +250bp from TSS) ChIP-seq signals at genes with the fastest Pol II decay versus other gene groups.

(G) Average distribution of IntS12 ChIP-seq signal, and violin plot of reads (-250bp to +250bp from TSS) at genes with the fastest Pol II decay versus those with more stable Pol II.

### Figure S2

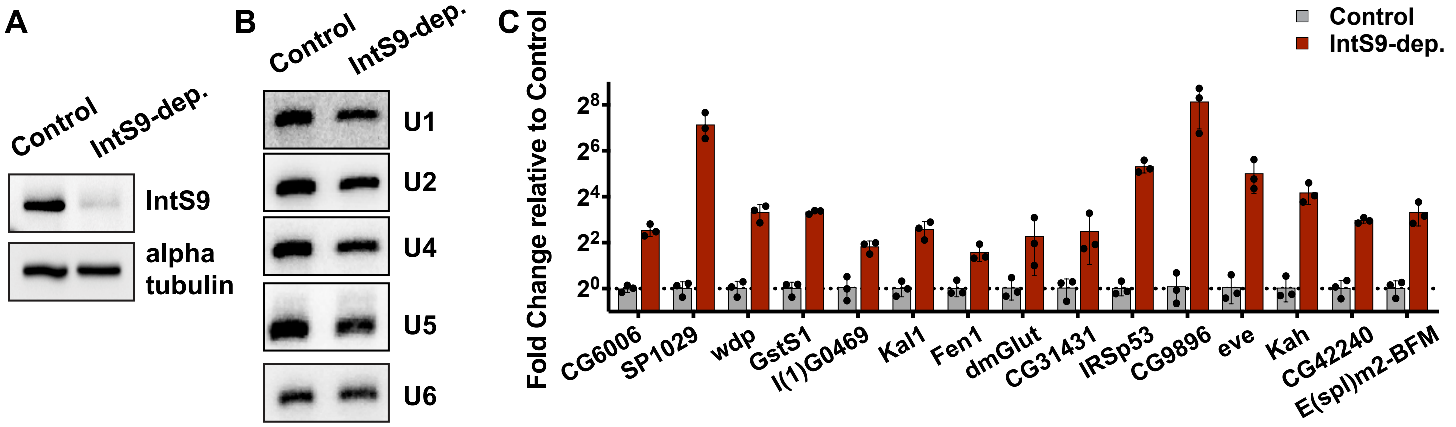


**D** Upregulated genes

| Top GO Molecular Function (Direct) | P-value |
|---|---|
| Gluthatione peroxidase activity | $1.5e^{-4}$ |
| Actin binding | $1.6e^{-4}$ |
| Oxidoreductase activity | $8.0e^{-4}$ |
| Calcium ion binding | $8.3e^{-4}$ |

| Top KEGG pathways | P-value |
|---|---|
| ECM-receptor interaction | $1.2e^{-6}$ |
| Glutathione metabolism | $1.1e^{-4}$ |
| Glycerolipid metabolism | $1.7e^{-3}$ |
| Metabolic pathways | $5.7e^{-3}$ |

| Top INTERPRO domains | P-value |
|---|---|
| EGF-like, conserved site | $1.3e^{-8}$ |
| Epidermal growth factor-like domain | $3.2e^{-8}$ |
| EGF-like calcium-binding | $7.7e^{-5}$ |
| ABC transporter, conserved site | $9.5e^{-5}$ |

**E** Downregulated genes

| Top GO Molecular Function (Direct) | P-value |
|---|---|
| Proton-transporting ATP synthase activity | $6.3e^{-4}$ |
| Cytochrome-c oxidase activity | $1.5e^{-3}$ |
| DNA binding | $6.5e^{-3}$ |
| Satellite DNA binding | $3.8e^{-2}$ |

| Top KEGG pathways | P-value |
|---|---|
| Oxidative phosphorylation | $7.0e^{-4}$ |
| Carbon metabolism | $2.0e^{-2}$ |
| Citrate cycle (TCA cycle) | $2.4e^{-2}$ |
| Valine, Leucine and isoleucine degradation | $7.4e^{-2}$ |

| Top INTERPRO domains | P-value |
|---|---|
| NOP5, N-terminal | $1.8e^{-2}$ |
| CDC6, C-terminal domain | $1.8e^{-2}$ |
| P-loop nucleoside triphosphate hydrolase | $1.9e^{-2}$ |
| Nop domain | $2.7e^{-2}$ |

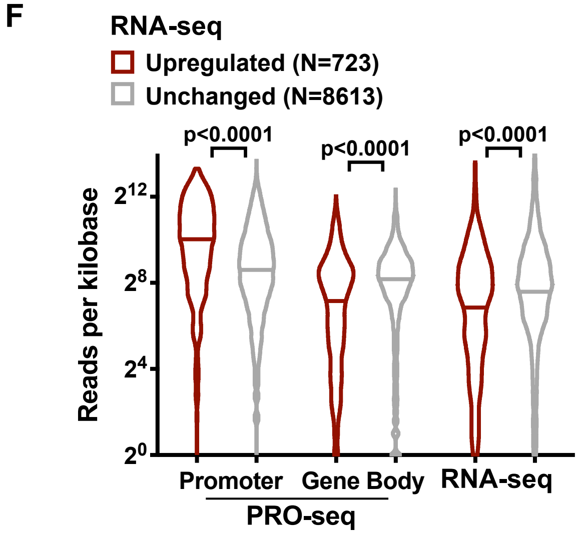


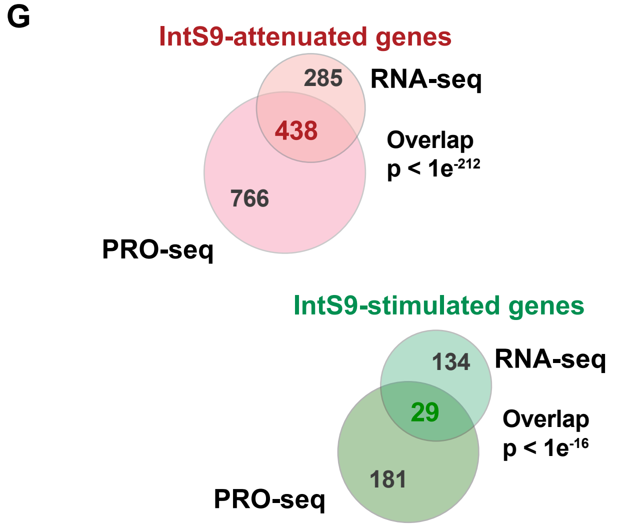


**Figure S2. Related to Figure 2. The Integrator complex attenuates expression of protein-coding genes.**
(A) Representative Western blot showing levels of IntS9 in cells following 60 h treatment with a control dsRNA, or a dsRNA targeting IntS9. Alpha-tubulin was used as a loading control.
(B) Representative Northern blots showing that mature snRNA levels are not affected upon IntS9-depletion. U6 snRNA biogenesis does not require the Integrator complex and serves as a loading control.
(C) RT-qPCR validation of representative genes affected by Integrator. mRNA levels from control and IntS9-depleted cells were normalized to RpS17 expression and the fold change upon IntS9 depletion is shown (mean ± SD, N=3).
(D-E) Gene Ontology analysis was performed on (D) the 723 transcripts upregulated or (E) the 163 transcripts downregulated by IntS9-depletion in the RNA-seq assay. This corresponded to 511 and 111 unique genes, respectively, that had defined Gene Ontology ID annotations in DAVID Tools (v6.8). The top four enriched functional categories, pathways and domains are reported.
(F) Shown are promoter (±150 nt from TSS) and gene body (+250 to +1250 nt from TSS) PRO-seq signal density at genes upregulated by IntS9-depletion, as compared to unchanged genes. Normalized RNA-seq counts are also shown. Violin plots depict range of values, with line indicating median. P-values are from Mann-Whitney test.
(G) Overlap between IntS9-affected genes in RNA-seq and PRO-seq assays. P-values were calculated using a hypergeometric test using a total of 9,499 active mRNA genes.

### Figure S3

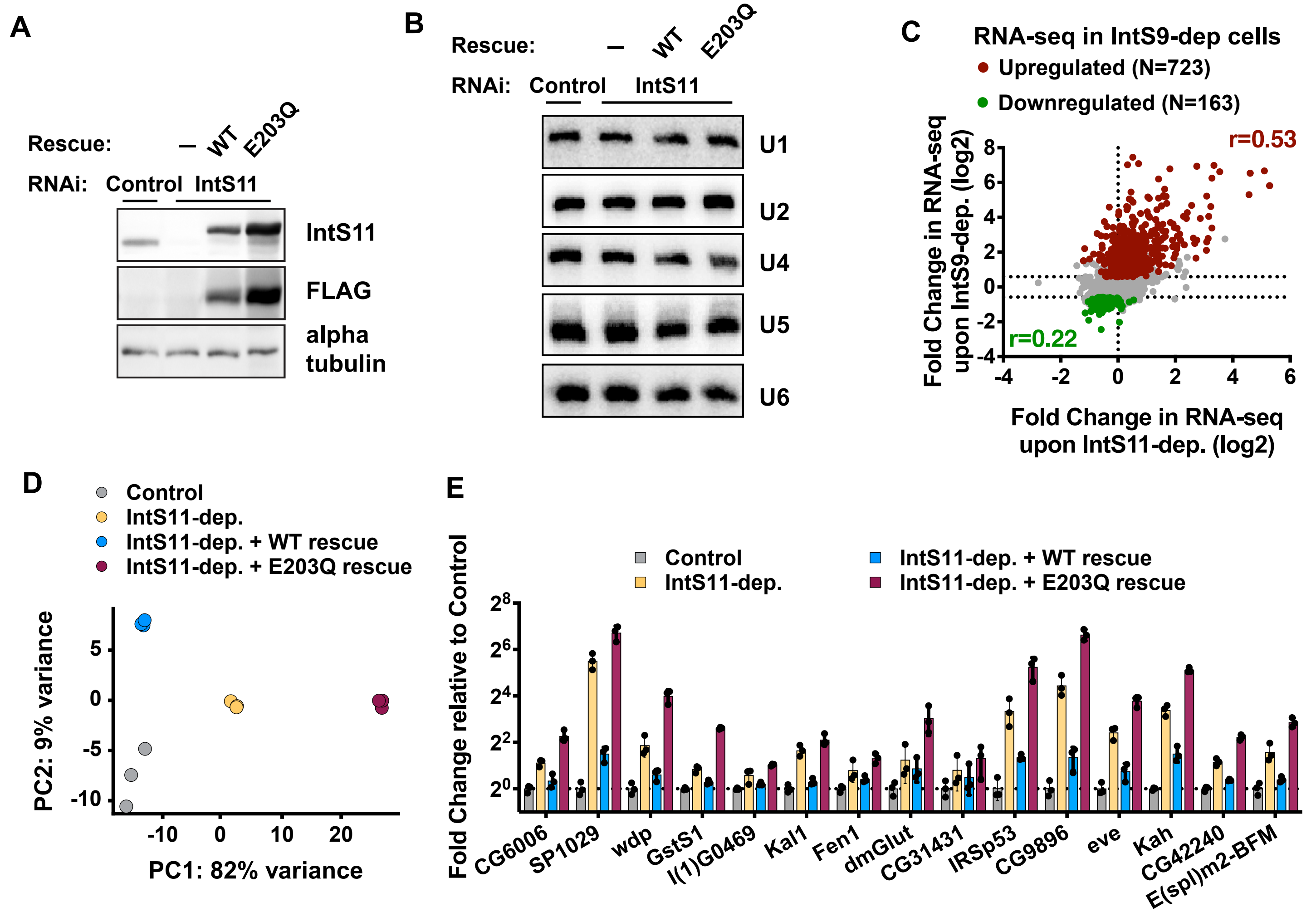


**Figure S3. Related to Figure 3. IntS11 catalytic activity is essential for attenuation of protein-coding genes.**

(A) Representative Western blot showing protein levels of endogenous IntS11 and the exogenous IntS11 WT and E203Q transgenes. Cells were treated with a control dsRNA or a dsRNA targeting IntS11 for 60 h. Alpha-tubulin was used as a loading control.

(B) Representative Northern blots showing that mature snRNA levels are not affected upon IntS11-depletion. U6 snRNA biogenesis does not require the Integrator complex and serves as a loading control.

(C) Comparison of the fold changes in steady-state RNA-seq signals upon IntS9 or IntS11-depletion is shown. Pearson correlations were calculated separately for upregulated and downregulated genes, indicating good agreement between signals for upregulated genes.

(D) 2-D principal component analysis of RNA-seq read counts at mRNA genes. Note that the Control cells and IntS11-depleted cells rescued with WT IntS11 are grouped together in Principle component 1 (PC1), which describes the majority (82%) of the variance among samples.

(E) RT-qPCR validation of representative genes supporting mRNA attenuation and rescue of expression by IntS11. Stably transfected cells were treated for 60h with a control dsRNA or dsRNA targeting IntS11. mRNA levels were normalized to RpS17 expression and the fold change is shown upon IntS11 depletion and the different rescue conditions (mean ± SD, N=3).

**Figure S4**

**A** **Upregulated genes**

| Top GO Molecular Function (Direct) | P-value |
|---|---|
| Oxidoreductase activity | $4.6e^{-5}$ |
| Gluthatione peroxidase activity | $9.6e^{-4}$ |
| Transcription factor activity, DNA binding | $1.7e^{-3}$ |
| Sterol-transporting ATPase activity | $3.4e^{-3}$ |

| Top KEGG pathways | P-value |
|---|---|
| Metabolic pathways | $2.8e^{-9}$ |
| Biosynthesis of antibiotics | $1.9e^{-8}$ |
| Fatty acid degradation | $3.3e^{-6}$ |
| ECM-receptor interaction | $3.0e^{-5}$ |

| Top INTERPRO domains | P-value |
|---|---|
| EGF-like, conserved site | $1.2e^{-9}$ |
| Epidermal growth factor-like domain | $1.7e^{-8}$ |
| NAD(P)-binding domain | $8.0e^{-6}$ |
| Fibronectin, Type III | $4.1e^{-5}$ |

**B** **Downregulated genes**

| Top GO Molecular Function (Direct) | P-value |
|---|---|
| Chromatin binding | $5.6e^{-8}$ |
| DNA binding | $1.3e^{-5}$ |
| POZ domain binding | $2.8e^{-4}$ |
| Protein binding | $9.7e^{-4}$ |

| Top KEGG pathways | P-value |
|---|---|
| Dorso-ventral axis formation | $6.8e^{-3}$ |

| Top INTERPRO domains | P-value |
|---|---|
| Immunoglobulin-like fold | $3.9e^{-5}$ |
| Fibronectin, Type III | $6.6e^{-4}$ |
| Immunoglobulin-like domain | $7.4e^{-3}$ |
| ENTH/ VHS | $2.2e^{-2}$ |

**C**

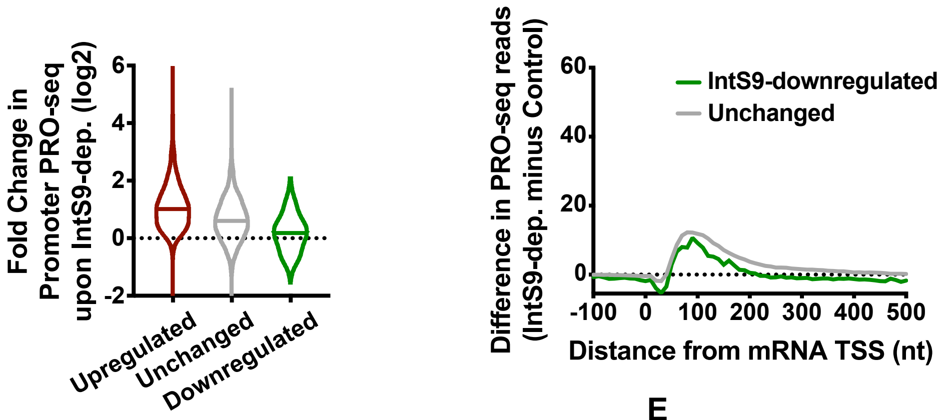


**D**

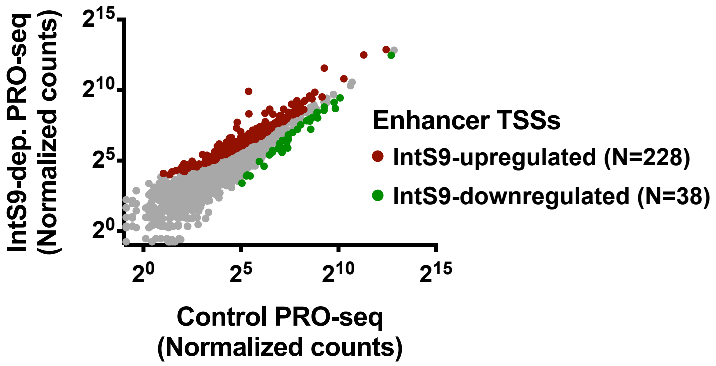


**E**

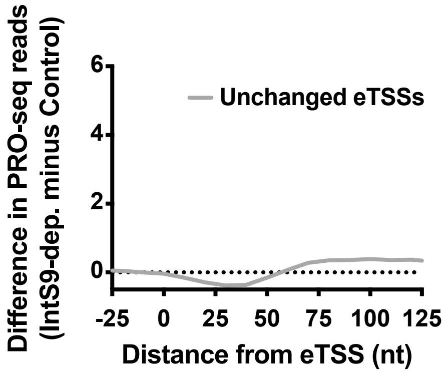


**Figure S4. Related to Figure 4. Loss of integrator complex causes transcription upregulation at protein-coding genes and enhancers.**

(A-B) Gene Ontology analysis of (A) the 1204 transcripts upregulated or (B) the 210 transcripts downregulated by IntS9-depletion, as defined from PRO-seq data. This corresponded to 1141 and 203 unique genes, respectively, that had defined Gene Ontology ID annotations in DAVID Tools (v6.8). The top four enriched functional categories, pathways and domains are reported. A single KEGG pathway was found to be enriched in downregulated genes.

(C) At left: Violin plots depict the change in promoter PRO-seq signal (± 150nt) upon IntS9-depletion for IntS9-affected genes (defined as in A) and unchanged genes (N=8085). Violin plots show range of values, with a line indicating median. At right: Difference in PRO-seq signal between IntS9-depleted and control cells is shown at downregulated (N=210) and unchanged genes (N=8085).

(D) Normalized PRO-seq signal at IntS9-upregulated enhancer RNA loci is shown downstream of enhancer TSSs. Upregulated eTSSs are defined from PRO-seq data from eTSS to +500bp, using $P<0.05$ and fold change >1.3.

(E) Difference in PRO-seq signal between IntS9-depleted and control cells is shown for enhancer loci that show no significant changes in PRO-seq signal upon IntS9-depletion (N=1232).

**Figure S5**

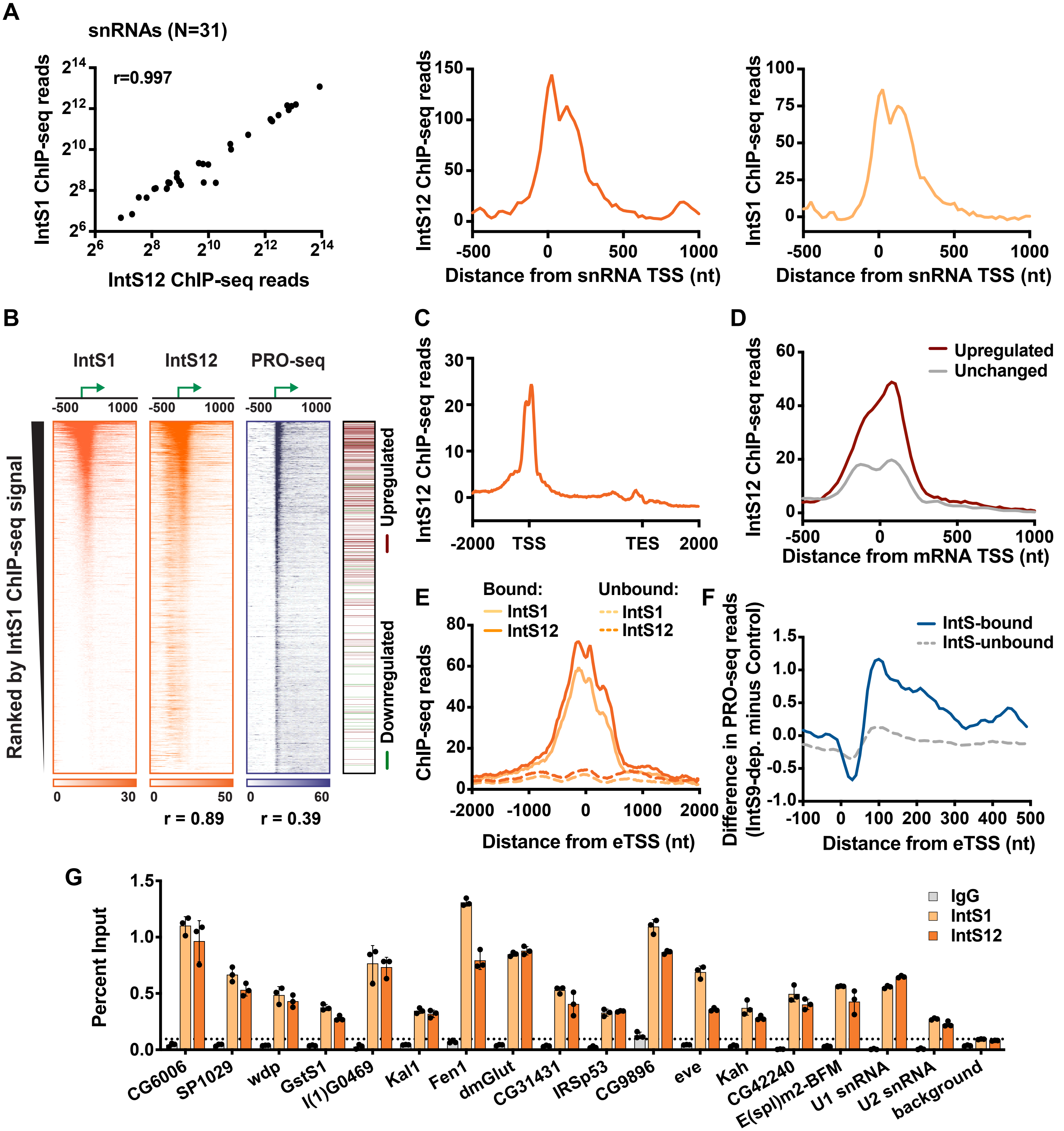


**Figure S5. Related to Figure 5. The Integrator complex is broadly associated with snRNA and mRNA genes.**
(A) Left: Signal from ChIP-seq for IntS1 versus IntS12 is shown at all 31 snRNA genes (using window from -150 to +350 bp from each TSS). Pearson correlation coefficient is indicated. Right: Average distribution of IntS12 and IntS1 ChIP-seq signal is depicted at snRNA loci. Read counts were summed in 25 nt bins, aligned on the TSS.
(B) Heatmap representations of IntS12 and IntS1 ChIP-seq signal and PRO-seq signal around mRNA TSSs (N=9499). Genes are ranked ordered by decreasing promoter-proximal IntS1 ChIP-seq signal (calculated from ±250 bp from TSS). Color bar at right indicates location of upregulated (red), downregulated (green) and unchanged (white) genes, based on PRO-seq in IntS9-depleted cells. Green arrow depicts TSS. Pearson correlation coefficients of promoter signals are shown below each heatmap.
(C) Distribution of IntS12 ChIP-seq signal along the transcription units of all active mRNA genes from TSS to transcription end site (TES), with bin size scaled by gene length.
(D) Average distribution of IntS12 ChIP-seq signal is shown at upregulated and unchanged genes.
(E) Average distribution of IntS1 and IntS12 ChIP-seq signal at IntS-bound eTSSs (N=691) vs. unbound (N=4182).
(F) Difference in PRO-seq signal between IntS9-depleted and control cells for IntS-bound and unbound eTSSs.
(G) ChIP-qPCR validation of IntS1 and IntS12 at promoters of representative Integrator target loci and snRNAs. ChIP signal is normalized relative to input (mean ± SD, N=3).

**Figure S6**

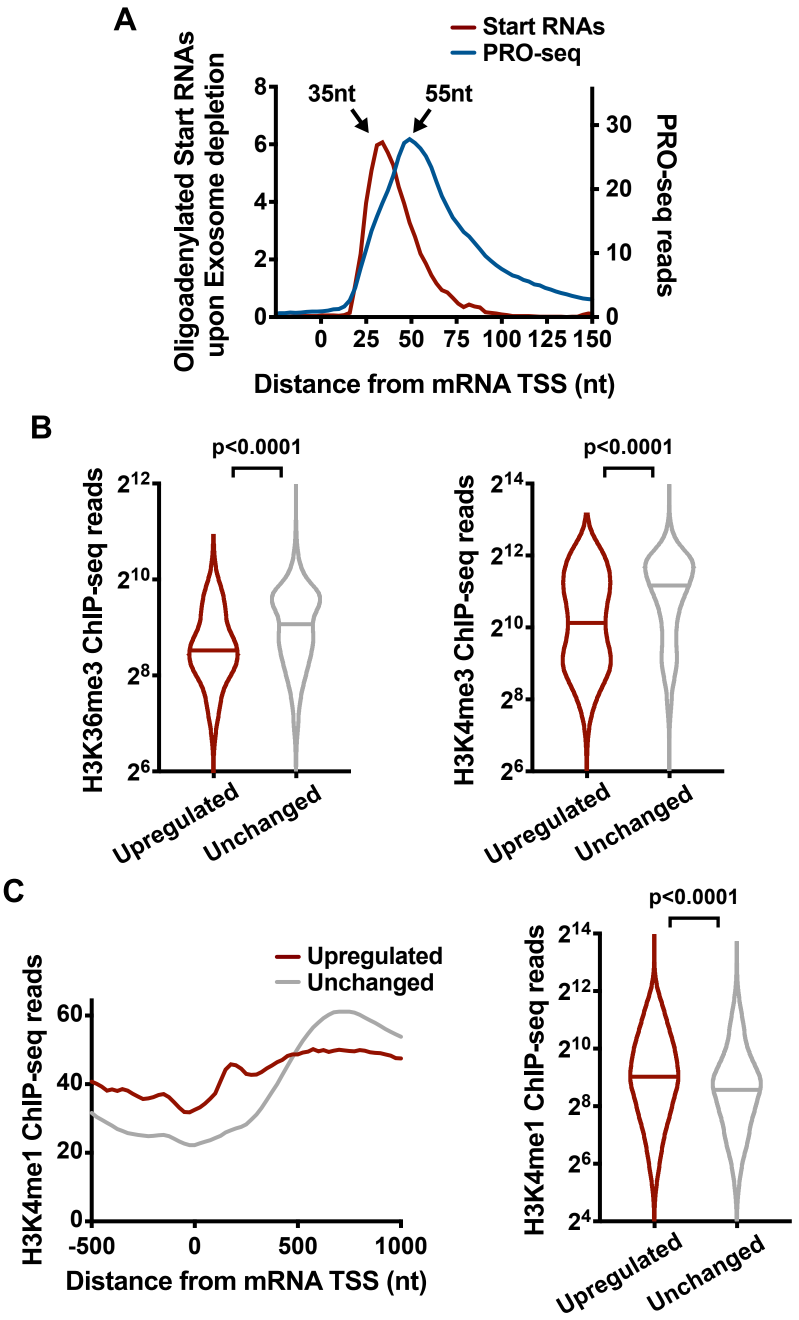


**Figure S6. Related to Figure 6. Integrator-repressed genes exhibit chromatin features consistent with unstable Pol II pausing and defective transcription elongation.**

(A) Average distribution of PRO-seq signal depicting the 3' ends of nascent RNA held within elongation complexes (blue), as compared to the 3' end locations of oligoadenylated RNAs identified in exosome-depleted cells (red). Shown are reads at genes upregulated by IntS9-depletion. The peak position of each distribution is indicated by an arrow.

(B) H3K36me3 (left) and H3K4me3 (right) ChIP-seq read counts are shown for genes upregulated or unchanged upon IntS9-depletion. Gene sets are defined as in Figure 4A. Windows used comprise TSS to +2000bp for H3K36me3, or TSS to +500bp from TSS for H3K4me3.

(C) At left: average distribution of H3K4me1 ChIP-seq signal is shown, aligned around TSSs, at upregulated or unchanged genes. At right: H3K4me1 ChIP-seq read counts (TSS to +500bp) are shown.

Figure S7

A

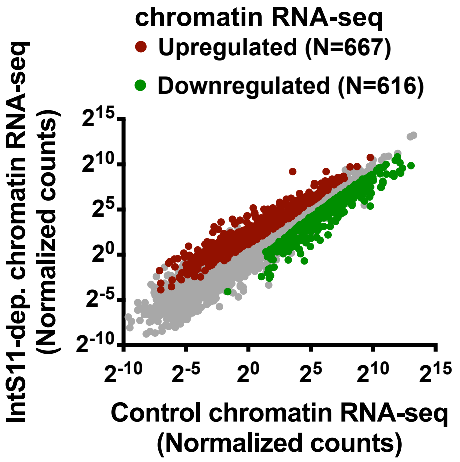


B **Upregulated genes**

| Top GO Molecular Function (Direct) | P-value |
|---|---|
| Calcium ion binding | $2.2e^{-7}$ |
| Extracellular matrix structural constituent | $2.4e^{-7}$ |
| Heparin binding | $1.5e^{-6}$ |
| Transmembrane receptor protein tyrosine phosphatase activity | $1.7e^{-4}$ |

| Top KEGG pathways | P-value |
|---|---|
| ECM-receptor interaction | $3.6e^{-11}$ |
| PI3K-Akt signaling pathway | $4.4e^{-9}$ |
| Focal Adhesion | $1.6e^{-7}$ |
| Pathways in cancer | $2.4e^{-5}$ |

| Top INTERPRO domains | P-value |
|---|---|
| Epidermal growth factor-like domain | $1.4e^{-8}$ |
| EGF-type aspartate/asparagine hydroxylation site | $3.5e^{-7}$ |
| EGF-like, conserved site | $7.8e^{-7}$ |
| EGF-like calcium-binding, conserved site | $1.3e^{-6}$ |

C **Downregulated genes**

| Top GO Molecular Function (Direct) | P-value |
|---|---|
| Protein binding | $1.4e^{-10}$ |
| poly(A) RNA binding | $3.3e^{-8}$ |
| ATP binding | $7.4e^{-8}$ |
| structural constituent of ribosome | $7.6e^{-8}$ |

| Top KEGG pathways | P-value |
|---|---|
| Cell cycle | $2.7e^{-11}$ |
| Ribosome | $6.9e^{-7}$ |
| Biosynthesis of amino acids | $6.6e^{-6}$ |
| DNA replication | $1.1e^{-5}$ |

| Top INTERPRO domains | P-value |
|---|---|
| Mini-chromosome maintenance | $1.2e^{-5}$ |
| Rossman-like alpha/beta/alpha sandwich fold | $3.3e^{-5}$ |
| Nuclei acid-binding, OB-fold | $9.4e^{-5}$ |
| Histone-fold | $1.1e^{-4}$ |

D

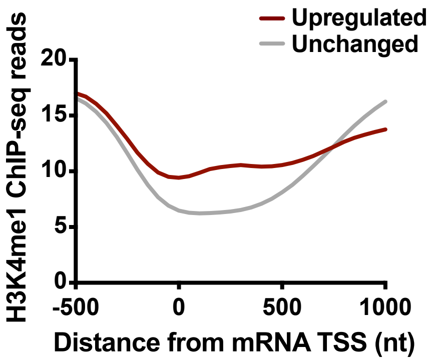


**Figure S7. Related to Figure 7. The Integrator complex affects expression of mammalian protein-coding genes under basal conditions.**

(A) Chromatin RNA-seq data from Lai *et al.,* 2015 were analyzed similarly to PRO-seq. Data are from HeLa cells depleted of IntS11 for 72h, as compared to cells expressing a scrambled shRNA. Depth normalized chromatin RNA-seq signal is shown. Significantly affected genes are defined using $P<0.0001$ and fold change >1.5.

(B-C) Gene Ontology analysis was performed on (B) the 667 genes upregulated or (C) the 616 genes downregulated by IntS11-depletion. This corresponded to 658 and 595 unique genes, respectively, that had defined Gene Ontology ID annotations in DAVID Tools (v6.8). The top four enriched functional categories, pathways and domains are reported.

(D) Average distribution of H3K4me1 ChIP-seq signal is shown, aligned around TSSs for upregulated or unchanged genes, defined as in A.
